# Tumor-infiltrating natural killer cell profiling for therapeutic stratification in patients with resectable non-small cell lung cancer

**DOI:** 10.64898/2026.01.01.696292

**Authors:** Atsushi Ito, Sung Wook Kang, Hee-Jin Jang, Ji Seon Shim, Claire Lee, Priyanka Ranchod, Gena Lee, Sonali Mitra, Jiyeon Jeon, Ayu H. Syarif, Yun Zhang, Jinyoung Byun, Younghun Han, Joshua Malo, Meera Patel, R. Taylor Ripley, Shawn S. Groth, Farrah Kheradmand, Bryan M. Burt, Francesca Polverino, Christopher I. Amos, Hyun-Sung Lee

## Abstract

**Objective:** Neoadjuvant chemoimmunotherapy has improved outcomes in resectable non-small cell lung cancer (NSCLC), yet its real-world implementation is often challenged by surgical delays, immune-mediated fibrosis, and postoperative complications. Smokers with NSCLC, despite a high risk for surgical morbidity, show enhanced responses to neoadjuvant chemoimmunotherapy. This study aimed to identify smoking-associated immune cell determinants that could guide treatment strategies.

**Methods:** Single-cell RNA sequencing (scRNAseq) was performed on 61 lung tissues from non-smokers and smokers to identify smoking-related immune compositions. The scRNAseq data from 19 invasive lung adenocarcinomas were used to validate their presence in the tumor-immune microenvironment. Bulk RNA sequencing data from 102 resected NSCLC and 24 NSCLC patients treated with neoadjuvant chemoimmunotherapy followed by surgery were used for *in silico* cellular deconvolution and outcome analyses.

**Results:** Among 135 lung cellular phenotypes, two natural killer (NK) cell subsets were strongly associated with smoking and chronic obstructive pulmonary disease (COPD) severity. “Stress-responsive” NK (NK_SR_) cells exhibited immature features and cytokine-responsive features, and “Adaptive and immunoregulatory NK” (NK_AIR_) cells showed mature features and elevated multiple immune checkpoint expression. High intratumoral NK_SR_ cells correlated with improved survival after surgery, particularly in current smokers. Conversely, tumors with low NK_SR_ and high NK_AIR_ cells responded more favorably to neoadjuvant chemoimmunotherapy.

**Conclusions:** Intratumoral NK cell phenotyping may aid in therapeutic stratification in patients with NSCLC. NK_SR_ cell preservation predicts benefit from upfront surgery, while NK_AIR_ cell enrichment indicates improved response to neoadjuvant chemoimmunotherapy. These NK cell profiles may help optimize treatment by balancing therapeutic benefit and risk.

**Graphic Abstract:** 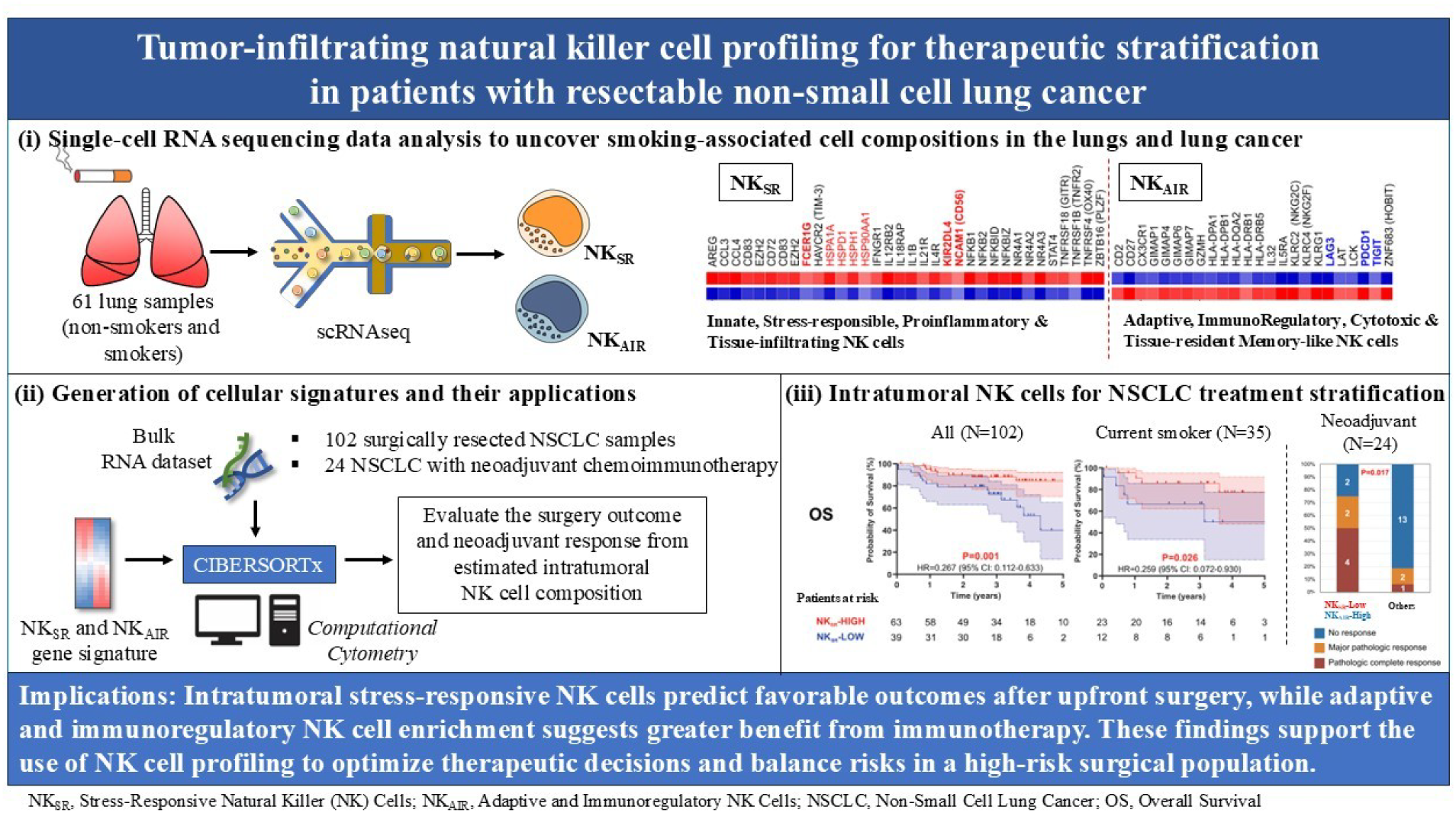

**Central Picture:** 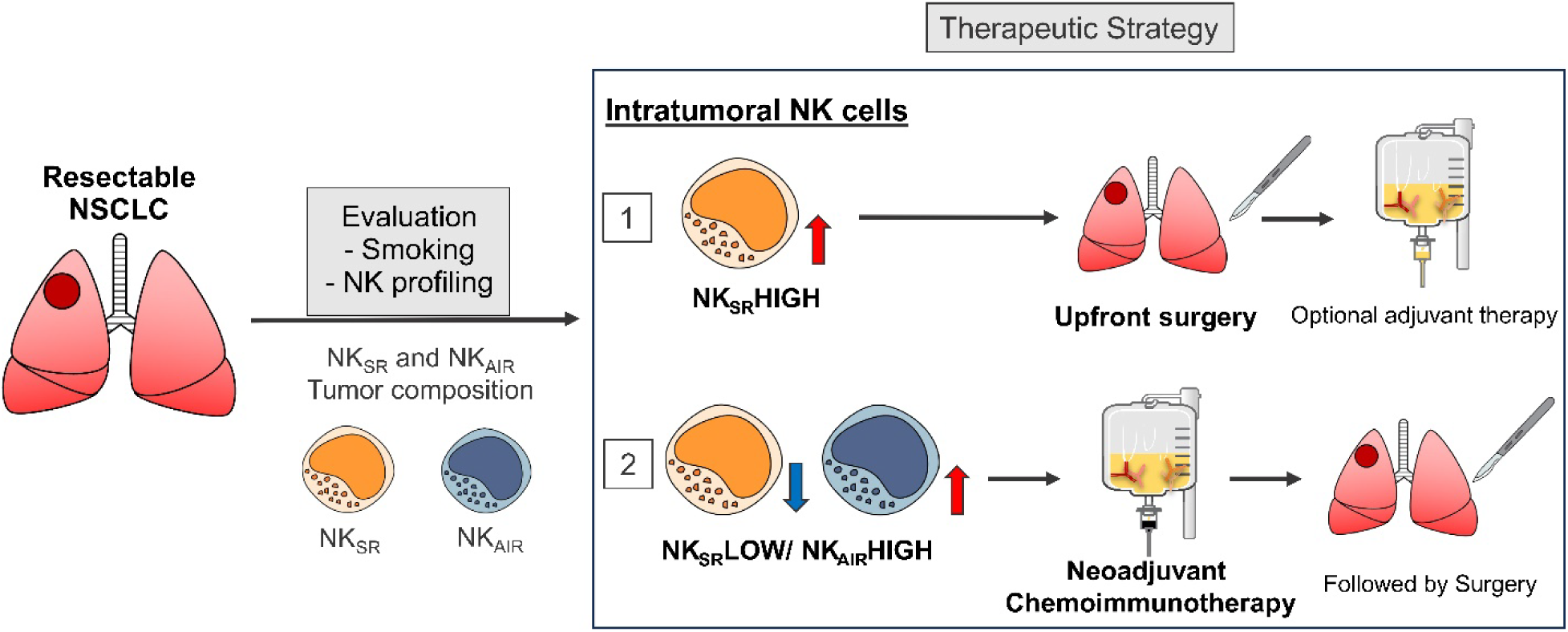

NK cell profiling for treatment stratification in resectable NSCLC

**Central Message:** Tumor-infiltrating NK cell profiling identifies distinct immune phenotypes that can inform personalized treatment strategies in resectable NSCLC.

**Perspective statement:** Abundance of intratumoral stress-responsive NK cells predicts favorable outcomes after upfront surgery, whereas enrichment of adaptive and immunoregulatory NK cells suggests greater benefit from neoadjuvant chemoimmunotherapy. These findings support incorporating NK-cell profiling into preoperative decision-making to optimize therapy selection and balance risks in the high-risk surgical population.

## Introduction

The introduction of immune checkpoint inhibitors (ICIs) has fundamentally reshaped the treatment landscape for advanced non-small cell lung cancer (NSCLC). Multiple predictive biomarkers, including tumor programmed cell death 1 ligand 1 (PD-L1) expression, tumor mutational burden (TMB), and smoking status, have become helpful indicators for clinical decision-making.^1–24^ As neoadjuvant immunotherapy or chemoimmunotherapy becomes increasingly adopted as a promising strategy for early-stage, resectable NSCLC,^25–29^ a clinical focus is shifting beyond palliative treatment for advanced disease toward curative-intent surgery, with maximizing long-term survival outcomes. Recent clinical trials, including CheckMate 816 and AEGEAN, have shown that neoadjuvant or perioperative chemoimmunotherapy significantly reduces the risk of relapse or death by 23% to 37% compared to chemotherapy alone.^25–27^ Notably, pathologic complete response (pCR) rates among current or former smokers receiving neoadjuvant chemoimmunotherapy ranged from 17% to 26%, whereas non-smokers achieved rates between 0% and 10%, likely attributable to their elevated TMB and increased tumor immunogenicity.^25–27^ However, the tumor-immune microenvironment (TiME) in early-stage NSCLC is biologically distinct from that of advanced disease,^30, 31^ limiting the direct applicability of biomarkers established in the metastatic setting. This discrepancy underscores an urgent need to identify biomarkers specifically tailored to the neoadjuvant setting in early-stage NSCLC.

From a surgical perspective, neoadjuvant chemoimmunotherapy presents several logistical and ethical challenges. Its real-world implementation often encounters multiple practical barriers, including surgical delays, cancellations, technical difficulties of surgery due to immune-mediated fibrotic changes, and increased postoperative complication rates.^32^ After neoadjuvant chemoimmunotherapy, 16% to 22% of patients did not undergo planned surgical resection due to disease progression, treatment-related adverse events, or new medical ineligibility.^25, 26, 28, 33^ Although smokers demonstrate superior responses to neoadjuvant chemoimmunotherapy, they also face higher risks of postoperative complications.^34–37^ This paradox highlights an unmet need for biomarkers that can stratify patients not only by therapeutic benefit but also by surgical risk.

Amid recent clinical advances, the identification and validation of reliable biomarkers in smokers is essential for optimizing neoadjuvant chemoimmunotherapy strategies and improving outcomes in patients with potentially resectable NSCLC. Therefore, we aimed to identify smoking-associated immune cell determinants by analyzing single-cell RNA sequencing (scRNAseq) data from the lung microenvironment of smokers and NSCLC tissues. Leveraging these scRNAseq-derived cellular signatures, we performed *in silico* deconvolution of bulk mRNA sequencing data to investigate the relationships between smoking-related immune cell compositions and clinical outcomes in patients with NSCLC who underwent either upfront surgical resection or neoadjuvant chemoimmunotherapy followed by surgery.

## Materials and Methods

### Study cohorts

This study was conducted under protocols approved by the Institutional Review Board at Baylor College of Medicine (BCM) (H-35782: approval date – 10/22/2021) and the University of Arizona (UA) (1811124026). Written informed consent was obtained from all participants for the collection of clinical data and biospecimens and the publication of the study data. At BCM, a prospectively maintained, single-institution database was reviewed retrospectively. The BCM cohort included 102 patients with histologically confirmed NSCLC who underwent complete pulmonary resection and mediastinal lymph node dissection. Staging was determined according to the 8^th^ edition of the TNM classification.^38^ At UA (Drs. Polverino and Malo), lung tissue samples were obtained from patients undergoing lung volume reduction surgery or lung transplantation for severe emphysema, or from patients undergoing lung resection for a solitary peripheral nodule, with sampled tissue taken from a region at least 10 cm away from the nodule. The UA cohort (N=61) included 13 non-smokers, 21 ever-smoker control subjects, 15 patients with mild chronic obstructive pulmonary disease (COPD)(GOLD stage 1-2), and 12 patients with severe COPD (GOLD stage 3-4). COPD was defined as a pre-bronchodilator forced expiratory volume in 1 second (FEV_1_)/forced vital capacity (FVC) ratio of less than 0.7 (70%). Disease severity was classified according to the GOLD criteria as follows: GOLD 1 (FEV1 ≥ 80%), GOLD 2 (50% ≤ FEV1 < 80%), GOLD 3 (30% ≤ FEV1 < 50%), and GOLD 4 (FEV1 < 30%).

### Single-cell isolation and scRNAseq

Lung tissues were collected for single-cell isolation as previously described.^30, 39^ In brief, tissues were minced and enzymatically digested using a human tissue dissociation kit (Miltenyi Biotec Inc., Auburn, CA, USA, Cat# 130-095-929). The digested mixture was sequentially filtered and washed twice. Then, cells were cryopreserved in a solution of 10% DMSO and 90% fetal bovine serum at −80°C. For long-term storage, cells were placed in the vapor phase of a liquid nitrogen storage system.

Single-cell suspensions were processed to isolate nuclei following the 10X Genomics protocol. Isolated nuclei were loaded into separate wells of a Chromium Next GEM Chip J (10X Genomics, Cat# 2000264) to generate gel beads-in-emulsion (GEMs), with each GEM containing uniquely barcoded beads for downstream single-cell capture and indexing. Nuclei concentration was adjusted to approximately 1,000 cells/µL to target recovery of up to 10,000 nuclei per sample. Sequencing libraries for simultaneous assay for transposase-accessible chromatin (ATAC) and 3’ gene expression were prepared using the Chromium Next GEM Single Cell Multiome GEM Kit A (10X Genomics, Cat# 1000282) according to the manufacturer’s protocol. All gene expression libraries were sequenced on the Illumina NovaSeq 6000 system using paired-end reads with dual indexing: 28 cycles for Read 1, 10 cycles for i7 Index, 10 cycles for i5 Index, and 90 cycles for Read 2. A sequencing depth of approximately 20,000 read pairs per cell was targeted. In this study, we used only gene expression data.

Sequenced reads were aligned to the human reference transcriptome (GRCh38) using the Cell Ranger (10x Genomics, v9.0.1)^40^ with default setting and generate gene-by-cell count matrices. Downstream quality control and analysis were performed in R using the Seurat package (v.5.3.0).^41^ Cells with fewer than 200 detected genes, greater than 20% mitochondrial gene content, or an abnormally high number of genes as doublets were excluded. Data normalization was performed using LogNormalize, with a scaling factor of 10,000. Dimensionality reduction was performed using uniform manifold approximation and projection (UMAP) for visualization.^42, 43^ Clustering was conducted using the Louvain algorithm with a resolution parameter set to 1.0.^44^ Initial cluster annotations were guided by differential gene expression analysis and confirmed using canonical marker genes to define major immune populations.

### RNA extraction, mRNA sequencing, and data analysis

Total RNA was extracted from fresh-frozen human NSCLC tissues using mirVana™ miRNA Isolation Kit (Thermo Fisher Scientific, Cat# AM1560). Ribosomal RNA was removed, and cDNA libraries were prepared using Illumina Stranded Total RNA Prep with Ribo-Zero Plus (Illumina, Cat# 20040529). Libraries were sequenced on the Illumina NovaSeq 6000 with 100 bp paired-end reads, generating a mean of 80 million reads per sample at the Human Genome Sequencing Core at BCM. Raw sequencing reads were processed according to established bioinformatic protocols. Initial quality assessment was performed using FastQC, followed by adapter trimming and quality filtering with Trimmomatic. The human reference genome (GRCh38) and corresponding annotation files were retrieved from Gencode (https://www.gencodegenes.org/) and utilized to generate the STAR aligner index. High-quality reads were subsequently aligned to the reference genome using STAR. Gene- and isoform-level quantification was performed with transcripts per million (TPM).

### Gene Set Enrichment Analysis

Gene set enrichment analysis (GSEA) was conducted using the GSEA 4.3.3 Desktop Application (http://www.gsea-msigdb.org/gsea/index.jsp).^45, 46^ Hallmark and BioCarta gene sets were obtained from the Molecular Signatures Database (MSigDB). This analysis involved 1000 random permutations for the gene sets and weighted enrichment statistics. The normalized enrichment score (NES) was used to assess enrichment, and significance was determined at a false discovery rate (FDR) < 0.05.

### Generation of scRNAseq-derived NK cell signature and quantification of NK cell Abundance

To generate a custom NK cell signature, we identified differentially expressed genes from our scRNA-seq dataset (P<10^-5^ with |fold change|>2). This signature was then incorporated into CIBERSORTx^47^ for digital cytometry analysis of bulk mRNA sequencing data. To infer the relative proportions of immune cell types, we utilized CIBERSORTx in relative mode, applying the LM22 reference matrix consisting of 22 immune cell phenotypes, including NK cells. Baseline NK cell composition in LM22 was replaced with new NK cell compositions based on scRNAseq-derived NK cell signature. Furthermore, to quantify absolute NK cell abundance, we employed CIBERSORTx in absolute mode, which calculates cell-type scores adjusted by the reference immune cell panel following quantile normalization. Unlike relative mode that reflects proportional composition, absolute mode provides a score representing the inferred abundance of each immune cell type, enabling more direct comparison across samples.

### Statistics

Cell compositions were expressed as percentages and summarized using means with standard deviations (SD). Pearson’s correlation analysis was performed to identify cell subsets positively or negatively associated with smoking status or COPD severity. Categorical variables were compared using Fisher’s exact test or Chi-square test, as appropriate. Continuous variables were analyzed using Student’s t-test for two-group comparisons and one-way ANOVA for comparisons involving more than two groups. For comparisons between tumors and their matched adjacent normal lung tissues, a paired t-test and Pearson correlation analysis were used. Survival analyses were conducted using the Kaplan-Meier method, and differences between groups were assessed using the log-rank test. Overall survival (OS) was defined as the time from surgery to death from any cause or last follow-up, while disease-free survival (DFS) was defined as the time from surgery to the first documented recurrence or death. To determine the cutoff value for stratifying patients by NK cell subset abundance, we employed a data-driven approach based on hazard ratios (HRs) using Cutoff Finder.^48^ Specifically, we plotted HRs across a range of potential cutoff values and selected the peak of the HR as the optimal threshold for survival analysis. Statistical significance was defined as P<0.05. Unless otherwise noted, all tests were two-sided. For comparisons of cell-type abundances between smokers and non-smokers in the limited scRNAseq datasets, one-sided tests were employed to assess the a priori hypothesis of higher abundance in smokers. All statistical analyses and graphic visualizations were performed using IBM SPSS 29.0 Statistics (IBM Corp) and GraphPad Prism 10.0 (GraphPad Software).

### Data Availability

The data generated in this study are available within the articles and its supplementary data files and next-generation sequencing data for all samples in this study have been deposited in the Gene Expression Omnibus accessions (GSE283245, GSE300685, and GSE302339). Additional information is available upon request to the corresponding author.

## RESULTS

### scRNAseq profiling revealed smoking-associated cellular alterations in the lung microenvironment

The study design is illustrated in **Figure 1**. Briefly, we identified cell populations associated with smoking status and COPD severity using lung scRNA-seq data (GSE302339). We then validated the presence of these smoking-related populations within the tumor-immune milieu using lung cancer scRNAseq datasets (GSE300685 and GSE131907^49^). To evaluate their clinical relevance, we derived cell-type signatures and applied them to NSCLC bulk RNA-seq cohorts via *in silico* cytometry (CIBERSORTx). Finally, we assessed the association between smoking-associated cell abundance and clinical outcomes in 102 completely resected lung cancers with 96 matched adjacent normal lungs (GSE283245) and examined response to neoadjuvant chemo-immunotherapy in an independent, publicly available cohort of 24 pre-treatment NSCLC tumors (GSE207422).

**Figure 1.**
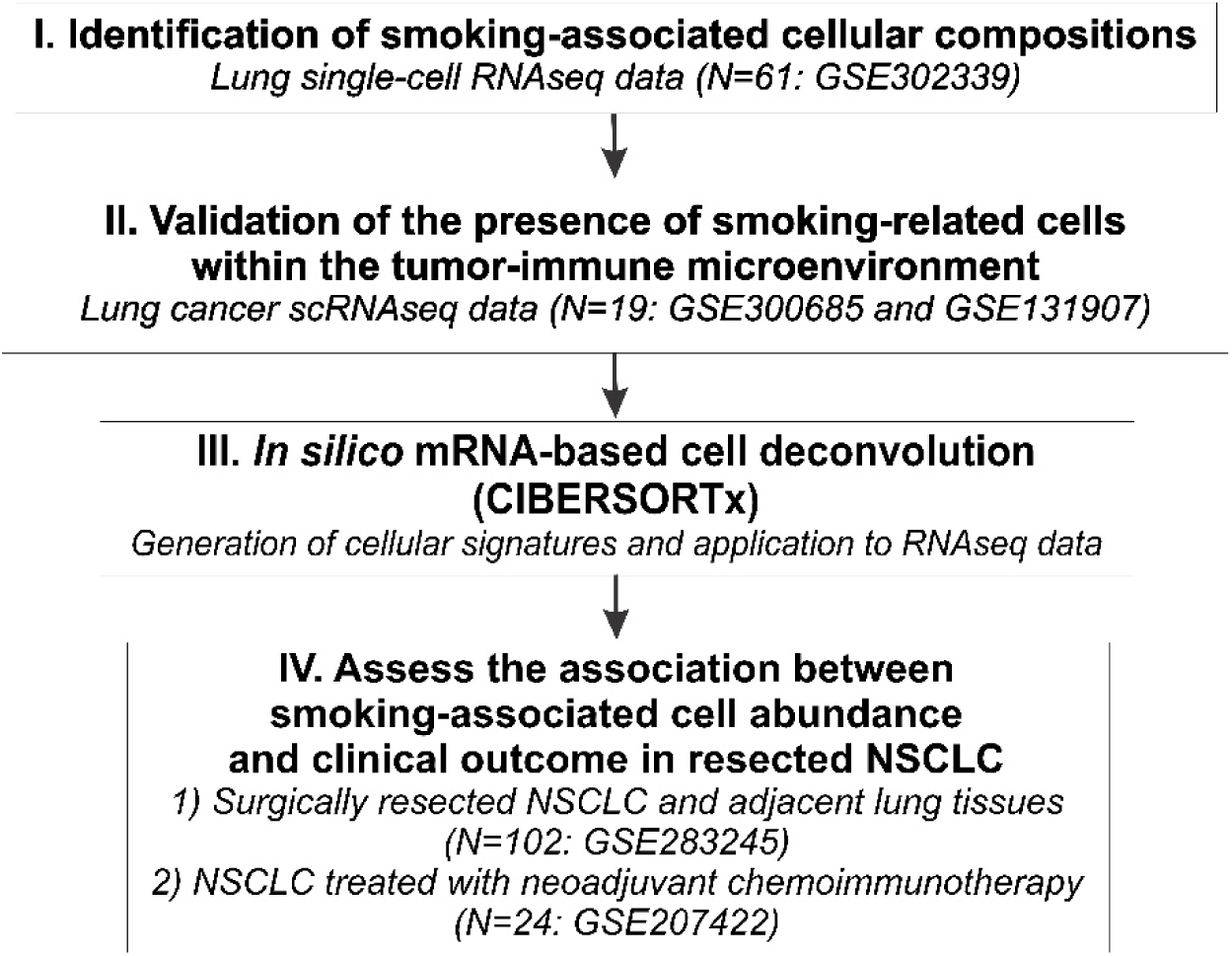
Study design overview. This study investigates smoking-associated immune cell determinants to predict clinical outcomes and guide treatment strategies in high-risk smokers with resectable NSCLC. First, we identified cell populations associated with smoking status and COPD severity using lung single-cell RNA-seq data (GSE302339). We then validated the presence of these smoking-related populations within the tumor immune microenvironment using lung cancer scRNA-seq datasets (GSE300685 and GSE131907). To evaluate their clinical relevance, we derived cell-type signatures and applied them to NSCLC bulk RNA-seq cohorts via *in silico* cytometry (CIBERSORTx). Finally, we assessed the association between smoking-associated cell abundance and clinical outcomes in 102 completely resected lung cancers with 96 matched adjacent normal lungs (GSE283245), and examined response to neoadjuvant chemo-immunotherapy in an independent, publicly available cohort of 24 pre-treatment NSCLC tumors (GSE207422). COPD, chronic obstructive pulmonary disease; NSCLC, non-small cell lung cancer.

We first analyzed lung scRNAseq data from a total of 61 patients, including 13 non-smokers, 21 smokers without COPD, 15 patients with mild COPD, and 12 patients with severe COPD, to identify the smoking-associated cellular alterations. The clinicopathological characteristics of this population are summarized in **Supplemental Table 1**. We analyzed 198,814 cells, which were classified into immune cells, including lymphoid and myeloid cells, and non-immune cells, including stromal, mesothelial, epithelial, and endothelial cells. The cellular composition was visualized using UMAP (**Figure 2A**), and individual patient-level distributions of major cell phenotypes were shown in **Figure 2B**. Comparisons across non-smokers, smokers, and patients with mild or severe COPD revealed no significant differences in the overall proportions of major cell types (**Figure 2C**).

**Figure 2.**
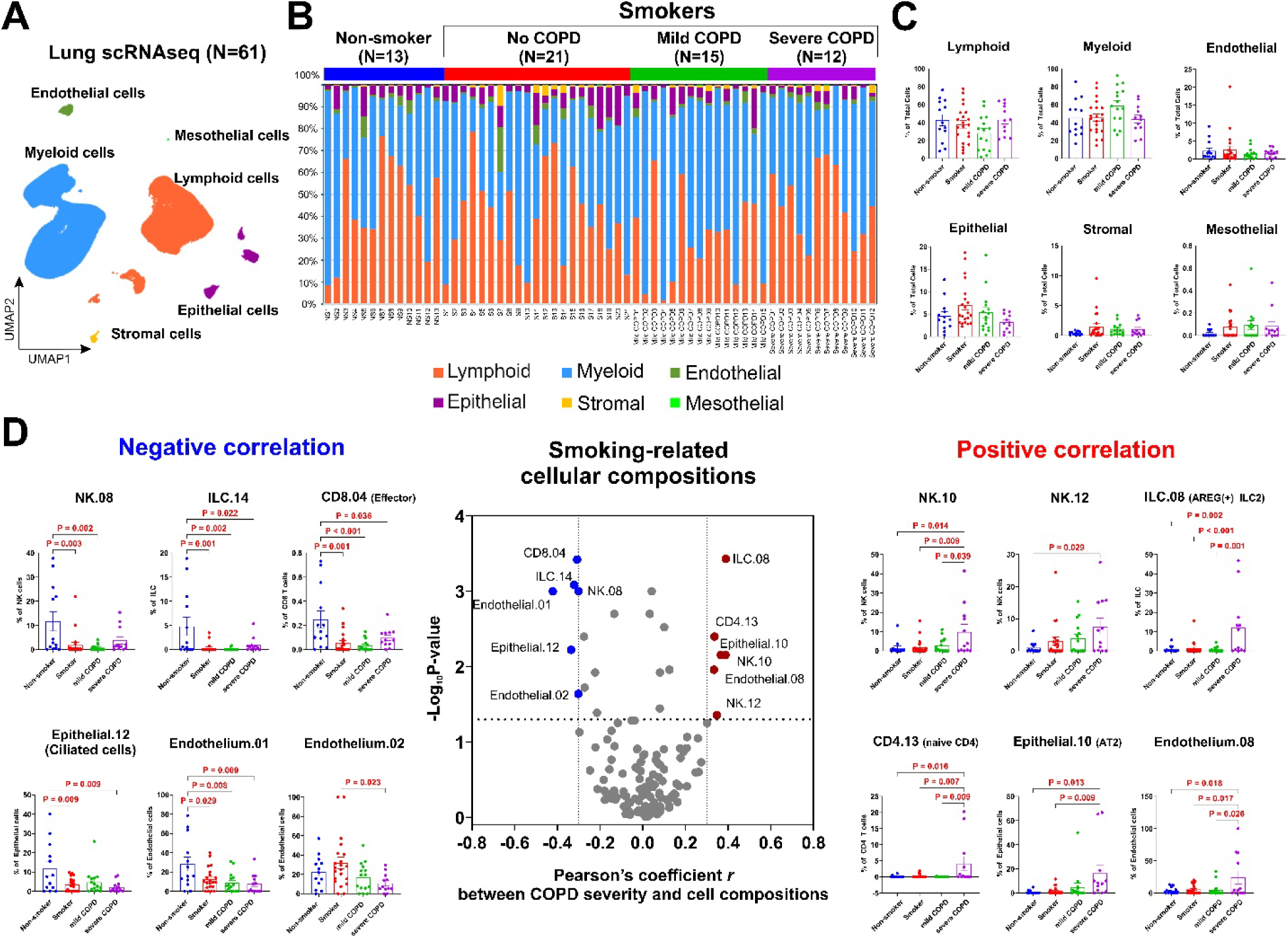
Single-cell transcriptomic profiling of lung immune and structural cell populations in non-smokers, smokers, and COPD patients. **A.** UMAP visualization of 198,814 single cells from lung tissues of 61 individuals (13 non-smokers, 21 smokers, 15 with mild COPD, and 12 with severe COPD), annotated into six major cell types: lymphoid, myeloid, epithelial, endothelial, stromal, and mesothelial cells. **B.** Stacked bar plot showing the proportional composition of the six major cell types across all individual samples grouped by clinical category: non-smoker, smoker, mild COPD, and severe COPD. **C.** Comparison of the proportion of each major cell type stratified by smoking status and COPD severity. Bars represent mean ± standard deviation (SD). **D.** Correlation of specific cell phenotypes with COPD severity. The central panel shows the volcano plot showing Pearson correlation coefficients and corresponding P-values between cell composition and COPD severity. Blue dots indicate significantly negatively correlated phenotypes (*r* ≤ –0.3, P<0.05); red dots indicate significantly positively correlated phenotypes (*r* ≥ 0.3, P<0.05). Left panels show negatively correlated cell types including NK.08, ILC.14, CD8.04 (effector), Epithelial.12 (ciliated), Endothelium.01, and Endothelium.02. Right panels show positively correlated cell types including NK.10, NK.12, ILC.08 (AREG⁺ ILC2), CD4.13 (naive CD4), Epithelial.10 (AT2), and Endothelium.08. Data are shown as mean ± SD. Statistical comparisons with 2-sided t-test were performed between groups. AT2, alveolar type 2 cells; COPD, chronic obstructive pulmonary disease; ILC, innate lymphoid cells; NK, natural killer cells; SD, standard deviations; UMAP, uniform manifold approximation and projection.

To gain deeper insight into smoking-induced cell alterations in the lung microenvironment, we extended our analysis with more detailed cell phenotypes. Lymphoid cells were classified into CD8 T cells, CD4 T cells, NK cells, B cells, and innate lymphoid cells (ILC), while myeloid cells were classified into macrophages, dendritic cells, monocytes, and neutrophils. Unsupervised clustering within each cell lineage revealed a total of 135 distinct cell phenotypes (**Supplementary Figure 1**). Correlation analysis was performed to identify cell phenotypes associated with smoking and COPD severity (**Figure 2D**). Six clusters demonstrated positive correlations with increasing COPD severity, including NK.10, NK.12, ILC.08 (AREG(+) ILC2), CD4.13 (Naïve CD4 T), Epithelial.10, and Endothelium.08. In contrast, six clusters, including NK.08, ILC.14, CD8.04 (Effector CD8 T), Epithelial.12 (ciliated cells), Endothelium.01, and Endothelium.02, showed negative correlations with COPD severity. Notably, among 12 significant clusters, three NK cell phenotypes, including NK.10, NK.12, and NK.08, exhibited significant associations with COPD severity. This finding suggests that NK cells may play a particularly pivotal role in COPD pathogenesis in the lung microenvironment. Given their strong association with COPD severity, NK cell populations were selected for in-depth analysis.

### Transcriptional and pathway analyses define distinct lung NK cell subsets

Building on the association of NK subsets with smoking status and COPD severity, we performed comprehensive transcriptional profiling of NK.10, NK.12, and NK.08 subpopulations to elucidate their functional properties (**Figure 3A**). UMAP analysis demonstrated that the combined NK.10 and NK.12 populations formed a distinct cluster separated from NK.08. Given the similarity of gene expression, NK.10 and NK.12 were collectively designated as NK subset 1, while NK.08 was classified as NK subset 2. Comparative transcriptomic analysis revealed markedly distinct gene expression profiles between NK cell subsets (**Figure 3B**). The NK subset 1 displayed a transcriptional signature characteristic of acute inflammatory activation, including upregulation of the tissue repair factor *AREG* and pro-inflammatory chemokines *CCL3, CCL4*. This subset demonstrated enhanced cytokine responsiveness through elevated expression of multiple cytokine signaling components responsive to IL-1, IL-12, IL-18, and IFN-γ (*IL1B*, *IL12RB2*, *IL18RAP*, *IL21R*, *IL4R*, *IFNGR1*), and a downstream signaling mediator STAT4. Coordinated upregulation of early and ligand-independent transcription factors (*NR4A1*, *NR4A2*, *NR4A3*) and NK-κB pathway components (*NFKB1*, *NFKB2*, *NFKBID*, *NFKBIZ*) indicated rapid transcriptional responses to inflammatory stimuli. Cellular stress adaptation was enhanced by increased expression of heat shock proteins (*HSPA1A*, *HSPD1*, *HSPH11*, *HSP90AA1*). This subset co-expressed the conventional NK marker (NCAM1/CD56), implying an immature phenotype,^50, 51^ as well as a high level of *ZBTB16* (*PLZF*), a transcription factor for invariant NKT (iNKT) cell development,^52, 53^ and *FCER1G*, which encodes a signaling adaptor that enhances both cytotoxicity and cytokine response.^54, 55^ These features collectively define a stress-responsive NK cell subset (NK_SR_) uniquely adapted to the inflammatory microenvironment of lungs damaged by smoking and capable of rapid effector responses.

**Figure 3.**
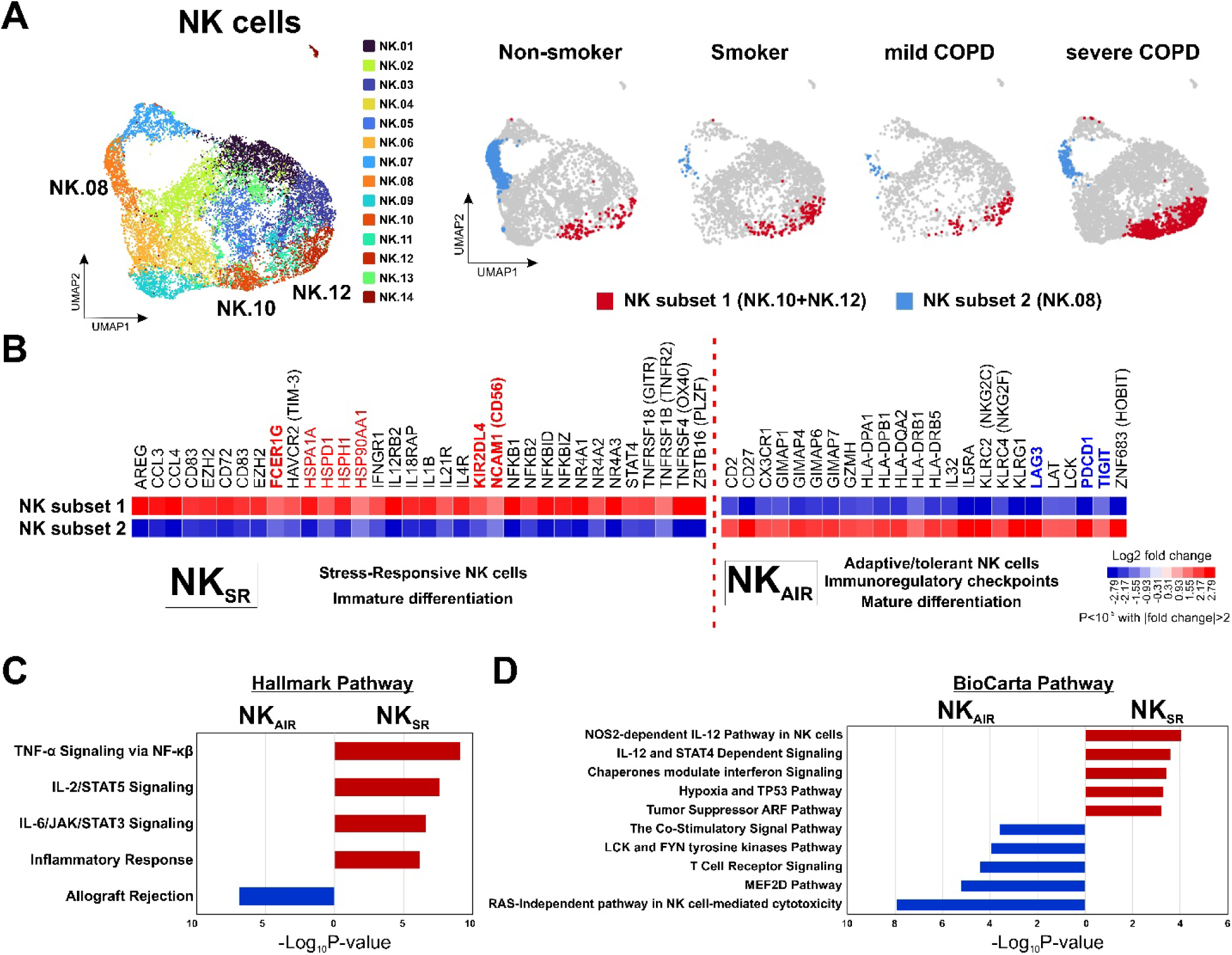
Characterization of smoking-associated NK cell phenotypes in the human lung. **A.** UMAP representation of NK cells derived from single-cell RNA-seq of 61 lung tissue samples, revealing 14 distinct NK cell clusters. NK.10 and NK.12 clusters (red dots) were enriched in smokers and COPD patients and designated as NK subset 1. In contrast, NK.08 (blue dots) was enriched in non-smokers and identified as NK subset 2. Distribution of NK subsets is shown across individuals stratified by smoking status and COPD severity. **B.** Differentially expressed gene profiling between NK subset 1 and subset 2. NK subset 1, characterized by elevated expression of CD56 (NCAM1), multiple heat shock protein genes, and cytokines, were defined as stress-responsive NK cells (NK_SR_). In contrast, NK subset 2 exhibited high expression of granzyme H and immune regulatory genes such as PDCD1, LAG3, and TIGIT, and were classified as adaptive and immunoregulatory NK cells (NK_AIR_). Genes with a log₂ fold change>|2| and P<10^-5^ are presented. **C.** Gene set enrichment analysis (GSEA) using MSigDB Hallmark pathways revealed that stress-responsive NK cells (NK_SR_) were enriched for pro-inflammatory signaling cascades, including TNF-α signaling via NF-κB, IL-2/STAT5, and IL-6/JAK/STAT3 pathways. In contrast, adaptive and immunoregulatory NK cells (NK_AIR_) showed enrichment for the allograft rejection pathway, indicative of immune regulatory activity. **D.** GSEA using BioCarta pathways further demonstrated that NK_SR_ cells were enriched for the nitric oxide synthase 2(NOS2)-dependent IL-12 signaling pathway, IL-12/STAT4 signaling, chaperone-modulated interferon signaling, as well as hypoxia and TP53-associated stress response pathways. In contrast, NK_AIR_ cells showed enrichment for pathways including RAS-independent NK cytotoxicity, LCK and FYN tyrosine kinase signaling, and co-stimulatory signaling, suggesting features of functional exhaustion yet retention of cytotoxic potential that may be amenable to reinvigoration. COPD, chronic obstructive pulmonary disease; GSEA, Gene set enrichment analysis; LAG3, lymphocyte Activating 3; NK, natural killer cells; PDCD1 Programmed Cell Death 1; TIGIT, T Cell Immunoreceptor With Ig And ITIM Domains; UMAP, uniform manifold approximation and projection.

In contrast, the NK subset 2 exhibited a transcriptional profile consistent with chronic activation and functional immune regulation (**Figure 3B**). This population was characterized by elevated expression of multiple immune checkpoint receptors (*PDCD1*, *LAG3*, and *TIGIT*)^56^, tissue residency markers, including *ZNF683* (*Hobit*)^57^ and *CX3CR1*^58, 59^, and upregulation of *CD27,*^60^ indicating long-term adaptation to the lung microenvironment with acquisition of a tissue-resident memory-like phenotype. Terminal differentiation was evidenced by upregulation of *KLRG1*^61^ and *KLRC2* (*NKG2C*)^62^. This subset displayed enhanced antigen presentation capabilities through upregulation of HLA class II genes (*HLA-DPA1*, *HLA-DPB1*, *HLA-DQA2*, *HLA-DRB1*, *HLA-DRB5*)^63, 64^ and T cell signaling components (*LAT*, *LCK*).^65, 66^ Increased expression of GTPase of immune-associated proteins (GIMAP) family members (*GIMAP1*, *GIMAP4*, *GIMAP7*), associated with lymphocyte survival and activation,^67^ along with the cytotoxic effector *GZMH* further supported a state of adaptive differentiation. These features define a population of tissue-resident NK cells shaped by chronic antigen exposure, with enhanced immunosurveillance capabilities but diminished effector function. We designated this subset as adaptive and immunoregulatory NK (NK_AIR_) cells, reflecting their putative role as immunoregulatory sentinels within chronically inflamed lungs.

Consistent with their transcriptional profiles, pathway enrichment analyses revealed that NK_SR_ cells were significantly enriched in pro-inflammatory signaling pathways, including TNF-α signaling via NF-κB, IL-2/STAT5 signaling, IL-6/JAK/STAT3 signaling, and the inflammatory response pathway (**Figure 3C**). NK_SR_ cells also demonstrated enhanced activity in IL-12-mediated signaling networks, including NOS2-dependent IL-12 signaling and IL-12/STAT4-dependent signaling pathway (**Figure 3D**). These pathway profiles indicate that NK_SR_ cells adopt a cytokine-driven activation mechanisms to facilitate robust pro-inflammatory responses and immune modulation. Conversely, NK_AIR_ cells demonstrated significant enrichment in allograft rejection and RAS-independent pathway in NK cell-mediated cytotoxicity, indicating preserved but possibly regulated cytolytic potential. Interestingly, NK_AIR_ cells exhibited enrichment in T cell receptor signaling, LCK and FYN tyrosine kinase pathway, and co-stimulatory signal pathway, indicating that they may retain cytotoxic potential.

### Distinct NK cell subsets are recapitulated in lung adenocarcinoma

To confirm the presence of NK subsets in the tumor immune microenvironment of lung cancer, we analyzed scRNAseq data from nineteen patients with invasive lung adenocarcinoma (LUAD) by integrating our 8 LUAD scRNAseq data (GSE300685) with publicly available 11 LUAD scRNAseq data (GSE131907: **Supplementary Figure 2A**).^49^ The smoking status and pathologic stage of these 19 patients are summarized in **Supplemental Table 2**. Intratumoral NK cells also had distinct clusters, whose differential gene expression patterns are highly consistent with findings from our previous transcriptomic analysis in **Figure 3B**. scRNAseq data confirmed the presence of both NK_SR_ and NK_AIR_ cell populations within the TiME **(Figure 4A**). NK_SR_ cells were enriched in tumors from smokers compared to non-smokers, consistent with smoking-associated expansion of this subset (**Figure 4B** and **4C**). Tumors from smokers also exhibited increased activated CD4 T (CD4 Tact) cells (**Figure 4D** and **4E**) and immunosuppressive M2-like tumor-associated macrophages (TAMs) (**Figure 4F**, **4G,** and **Supplementary Figure 2B**). The concurrent enrichment of cytokine-releasing NK_SR_ cells with activated lymphocytes, and M2-like TAMs suggests a transitional state - poised between acute immune activation and emerging immune evasion.

**Figure 4.**
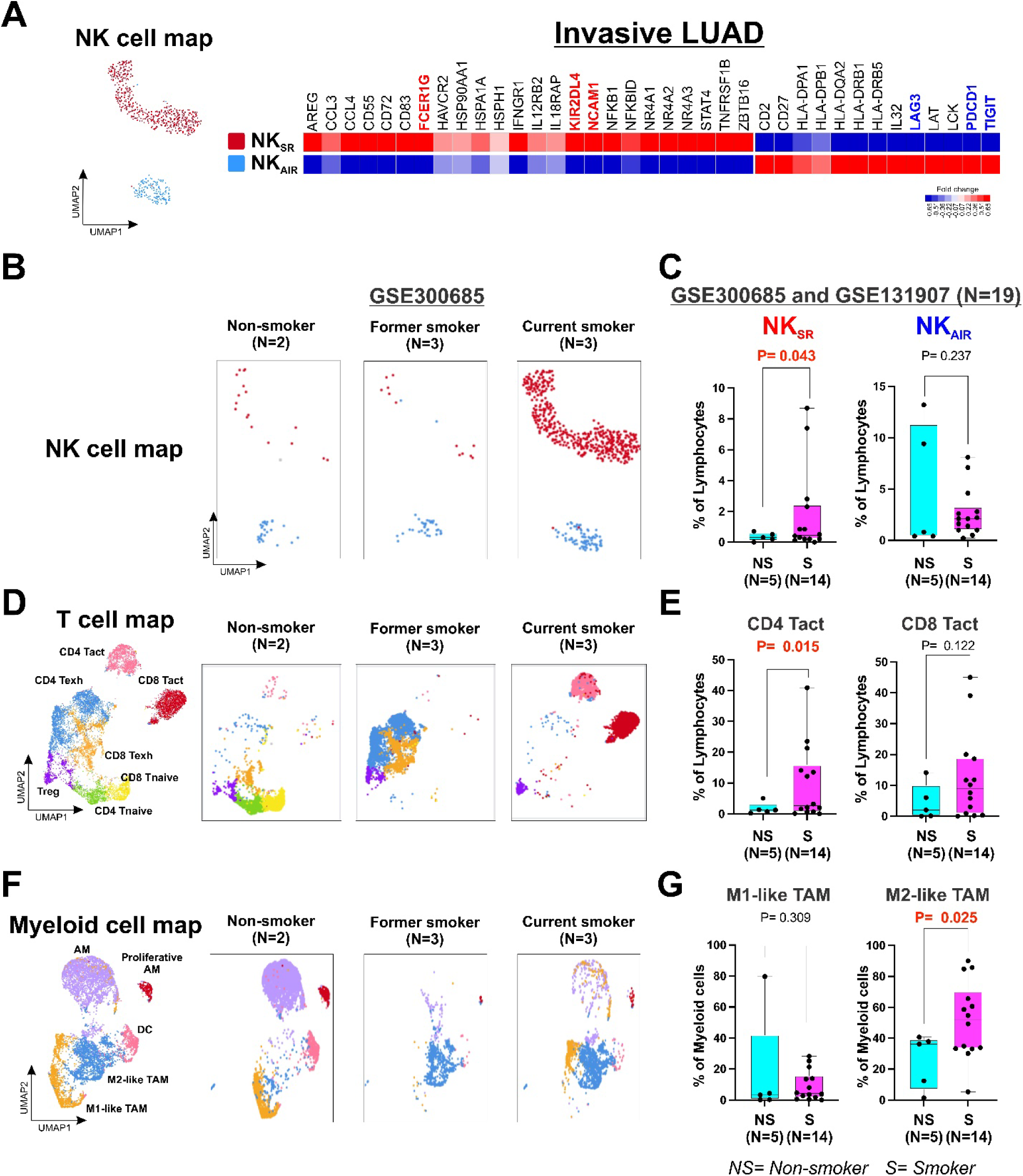
Distinct NK cell subsets are recapitulated in lung adenocarcinoma. **A.** UMAP visualization of NK cells from invasive lung adenocarcinoma (LUAD) reveals two distinct subsets: stress-responsive NK cells (NK_SR_) and adaptive/immunoregulatory NK cells (NK_AIR_). The heatmap displays differentially expressed genes between these subsets. NK_SR_ cells upregulate stress-related genes and inflammatory cytokines, whereas NK_AIR_ cells express high levels of immunoregulatory markers. These profiles are consistent with the NK phenotypes identified in **Figure 3B**. **B.** UMAPs depict the distribution of NK_SR_ and NK_AIR_ cells across three clinical subgroups: non-smokers (N=2), former smokers (N=3), and current smokers (N=3). Despite limited NK-cell counts, NK_SR_ cells were highly enriched in tumors from current smokers, compared to non-smokers and former smokers, **C.** Integrated scRNA-seq analysis (N=19) demonstrates significantly higher NK_SR_ cell proportions in smokers versus non-smokers, with no difference in NK_AIR_. Data are shown as mean ± SD. **D.** UMAPs of T cell subsets across the same clinical groups identify naïve CD4 T cells, activated CD4 T cells (Tact), exhausted CD4 T cells (Texh), regulatory T cells (Treg), naïve CD8 T cells, CD8 Tact, and CD8 Texh cells. **E.** In the integrated LUAD cohort (N=19), intratumoral activated CD4 T cells are significantly enriched in smokers. Other T cell populations had no differences between non-smokers and smokers (**Supplementary Figure 2**). **F.** UMAPs of myeloid cell populations show alveolar macrophages (AM), proliferative AMs, dendritic cells (DCs), M1-like tumor-associated macrophages (TAMs), and M2-like TAMs. **G.** Tumors from smokers were enriched for immunosuppressive M2-like TAMs. Other myeloid cell populations had no differences between non-smokers and smokers (**Supplementary Figure 2**).

### NK cell-based subtyping stratifies treatment outcomes in resectable NSCLC

To investigate the infiltration patterns of defined NK cell subsets between tumor and adjacent normal tissues in patients with NSCLC, we derived scRNAseq-based NK cell signatures to identify NK_SR_ and NK_AIR_ from bulk RNA sequencing data (**Supplementary Table 3**). Lung scRNAseq data showed that the average gene expression of these NK signatures was specific to the corresponding NK_SR_ and NK_AIR_ populations (**Supplemental Figure 3**). We then applied these NK signatures in CIBERSORTx digital cytometry^47^ across three datasets: 102 treatment-naïve NSCLC tumor samples (GSE283245), 96 matched adjacent normal lung samples (GSE283245), and publicly available 24 pre-treatment NSCLC tumors from patients who received neoadjuvant chemoimmunotherapy (GSE207422).^68^ The clinicopathological characteristics of our NSCLC cohort are summarized in **Table 1**.

**Table 1.** Patient characteristics.

| **Variables** | **All Patient** |
| --- | --- |
| **Number of patients** | 102 |
| **Age, median (range)** |  |
| **Sex, n (%)** |  |
| Male | 53 (51.9) |
| Female | 49 (48.0) |
| **Race, n (%)** |  |
| White | 72 (70.6) |
| Black | 18 (17.6) |
| Asian | 6 (5.9) |
| Unknown | 6 (5.9) |
| **Smoking status, n(%)** |  |
| Never | 26 (25.5) |
| Former | 41 (40.2) |
| Current | 35 (34.3) |
| **FEV1/FVC** |  |
| **Image finding** |  |
| Tumor size, cm (range) | 2.7 (1 - 7.5) |
| SUVmax (range) | 4.9 (0.8 - 31.3) |
| **Histology, n (%)** |  |
| Adenocarcinoma | 77 (75.5) |
| Squamous cell carcinoma | 25 (24.5) |
| **Grade, n (%)** |  |
| Well differentiated | 7 (6.9) |
| Moderately differentiated | 72 (70.6) |
| Poorly differentiated | 18 (17.6) |
| Unknown | 5 (4.9) |
| **Pathological Stage, n (%)** |  |
| IA | 44 (43.1) |
| IB | 22 (21.6) |
| IIA | 3 (2.9) |
| IIB | 18 (17.6) |
| IIIA | 15 (14.7) |

*FEV1*, forced expiratory volume in 1 second; *FVC*, forced vital capacity; *NSCLC*, non-small cell lung cancer; *SUVmax*, maximum standardized uptake value.

Immune deconvolution revealed a significant reduction of NK_SR_ cells in tumors compared to matched adjacent normal lung tissues (P=5.0×10^-5^ by paired t-test). NK_AIR_ cell levels showed no significant difference between tumor and normal tissues (**Figure 5A**). While NK_AIR_ cell abundance did not correlate between tumor and normal tissues, tumor NK_SR_ levels showed a positive correlation with NK_SR_ in matched adjacent normal tissues (**Figure 5B**), indicating that host/tissue context may partly shape the NK_SR_ phenotype within the tumor. Stratification by smoking status (current smokers [CS], former smokers [FS], and non-smokers [NS]) showed that although NK_SR_ cells are significantly enriched in the smoker lung microenvironment, tumor-specific exclusion of NK_SR_ cells was most pronounced in current smokers (P=5.3×10⁻⁴ by paired t-test) and modestly observed in non-smokers (P=0.034 by paired t-test) (**Figure 5C**). Together, these findings suggest smoking-associated modulation of NK-cell infiltration and selective evasion of NK_SR_ cells during tumorigenesis.

**Figure 5.**
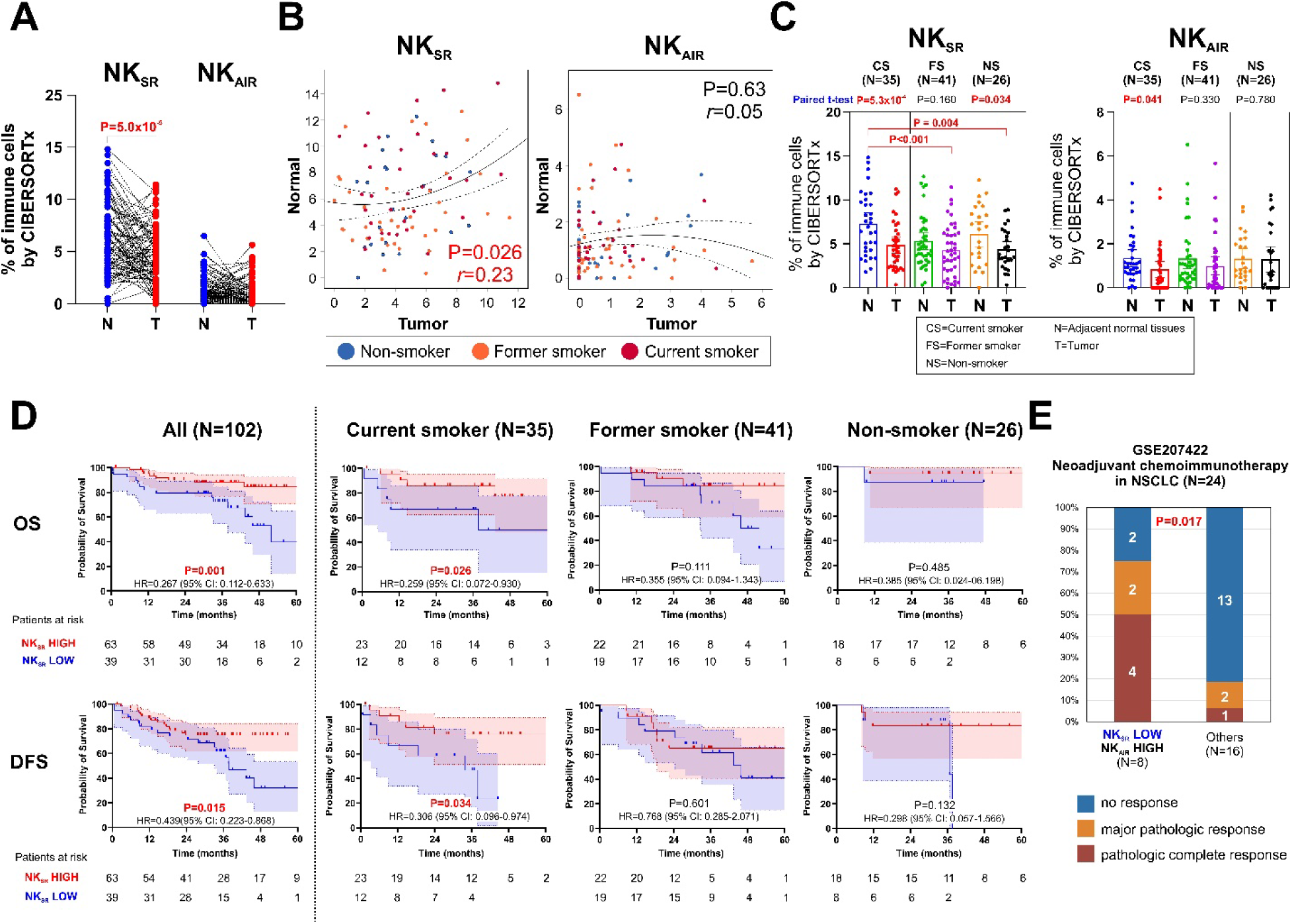
NK cell-based subtyping stratifies treatment outcomes in resectable NSCLC. **A.** Immune deconvolution revealed a significant reduction of NK_SR_ cells in tumors compared to matched normal tissues (P=5.0×10^-5^ by paired t-test). NK_AIR_ cell levels showed no significant difference between tumor and normal tissues. **B.** NK_SR_ abundance in tumors positively correlates with that in adjacent normal tissues, whereas NK_AIR_ shows no such correlation. **C.** Stratification by smoking status (current smokers [CS], former smokers [FS], non-smokers [NS]) showed that NK_SR_ cell exclusion from tumors was most pronounced in current smokers (P=5.3×10^-4^ by paired t-test) and modestly observed in non-smokers (P=0.034 by paired t-test). NK_AIR_ cells were significantly reduced in tumors only in current smokers (P=0.041 by paired t-test), suggesting smoking-associated modulation of NK cell infiltration and selective evasion of NK_SR_ cells during tumorigenesis. **D.** Kaplan–Meier survival analyses of overall survival (OS) and disease-free survival (DFS) in 102 resected NSCLC patients stratified by intratumoral NK_SR_ cell abundance. Patients with high NK_SR_ infiltration had significantly improved OS (P=0.001, HR=0.267) and DFS (P=0.015, HR=0.439). These survival benefits were especially evident in current smokers, while no significant survival difference was seen in non-smokers or former smokers. **E.** Analysis of a publicly available dataset (GSE207422) comprising 24 NSCLC patients treated with neoadjuvant chemoimmunotherapy. Patients were grouped by intratumoral NK_SR_ and NK_AIR_ abundance. The group with low NK_SR_ and high NK_AIR_ had significantly greater treatment responses, including higher rates of pathologic complete response (pCR) and major pathologic response (MPR) (P = 0.017), compared to all other NK cell profile combinations.

Given the more pronounced distributional differences in NK_SR_ cells between tumors and adjacent normal tissues, we next assessed their prognostic relevance. The 102 patients were classified into NK_SR_ HIGH and NK_SR_ LOW groups based on absolute NK_SR_ cell abundance score, using a cutoff value of 1.83 (**Supplementary Figure 4A**). The NK_SR_ HIGH group demonstrated significantly better OS (P=0.001, HR=0.267, 95% CI: 0.112-0.633) and DFS (P=0.015, HR=0.439, 95% CI: 0.223-0.868) (**Figure 5D**). Interestingly, this survival benefit of NK_SR_ HIGH group was particularly evident in current smokers, with significantly prolonged OS (P=0.026, HR=0.259, 95% CI:0.072-0.930) and DFS (P=0.034, HR=0.306, 95% CI: 0.096-0.974) (**Figure 5D**). In contrast, NK_AIR_ abundance did not correlate with survival outcomes (**Supplementary Figure 4B**).

Univariable Cox regression analysis to assess the prognostic impact of intratumoral NK_SR_ cell abundance showed that female sex, non-smoking status, lower cumulative tobacco exposure (pack-year), LUAD histology, earlier pathologic stage, and higher NK_SR_ cell abundance in the tumor were all significantly associated with better OS (**Supplementary Table 4**). In the multivariable analysis incorporating these factors, higher NK_SR_ cell abundance in the tumor remained independently associated with favorable outcomes, including better OS (*P* = 0.019; HR, 0.348; 95% CI, 0.144-0.841) and DFS (*P* = 0.029; HR, 0.459; 95% CI, 0.228-0.923). Advanced pathological stages (stage II/III) independently predicted worse DFS (*P* = 0.047; HR, 2.066; 95% CI, 1.010-4.226) (**Table 2**).

**Table 2.** Multivariable regression analysis of survival.

| Variables | Overall survival |  |  | Disease-free survival |  |  |
| --- | --- | --- | --- | --- | --- | --- |
|  | P values | HR | 95% CI | P values | HR | 95% CI |
| Sex<br>(female vs. male) | 0.134 | 0.450 | 0.159-1.280 | 0.624 | 0.828 | 0.390-1.759 |
| Smoking status<br>(Smoker vs. Non-smoker) | 0.220 | 2.570 | 0.571-11.595 | 0.341 | 1.573 | 0.620-3.989 |
| Histology<br>(LUSC vs. LUAD) | 0.181 | 1.825 | 0.755-4.410 | 0.358 | 1.418 | 0.674-2.984 |
| Pathologic stage<br>(II/III vs. I) | 0.287 | 1.622 | 0.666-3.947 | **0.047** | 2.066 | 1.010-4.226 |
| High NK<sub>SR</sub><br>in the tumor | **0.019** | 0.348 | 0.144-0.841 | **0.029** | 0.459 | 0.228-0.923 |

CI, confidence interval; HR, hazard ratio; LUSC, lung squamous cell carcinoma; LUAD, lung adenocarcinoma; NK<sub>SR</sub>, stress-responsive NK cells.

### Tumor-infiltrating NK cell composition predicts response to neoadjuvant chemoimmunotherapy in NSCLC

Given that smokers appear to benefit from neoadjuvant chemoimmunotherapy,^25–27^ NK_SR_-absent tumor is associated with unfavorable survival, and NK_AIR_ cells exhibit immunoregulatory signals, we hypothesized that the intratumoral balance of these NK cell subsets may influence the efficacy of neoadjuvant chemoimmunotherapy. To test this hypothesis, we analyzed mRNA data from 24 NSCLC patients who underwent complete pulmonary resection following neoadjuvant chemoimmunotherapy with PD-1 blockade and chemotherapy (GSE207422).^68^ Patients were grouped by intratumoral NK_SR_ and NK_AIR_ abundance score. The group with NK_SR_ LOW and NK_AIR_ HIGH had significantly greater treatment responses, including higher rates of pCR and major pathologic response (MPR) (P=0.017), compared to all other NK cell profile combinations (**Figure 5E**). Specifically, 4 of 8 (50%) in this group achieved a pCR, and an additional 2 patients (25%) exhibited an MPR, resulting in a combined pCR/MPR rate of 75%. In contrast, among patients with other NK cell profile combinations, only 1 of 16 (6.3%) achieved pCR and 2 of 16 (12.5%) exhibited MPR. These findings suggest that tumors with high NK_AIR_ and low NK_SR_ cell abundance are more likely to respond to neoadjuvant chemoimmunotherapy, potentially serving as a predictive biomarker for treatment stratification.

## DISCUSSIONS

In this study, we identified two transcriptionally and functionally distinct NK cell subsets, stress-responsive NK cells and adaptive/immunoregulatory NK cells. These NK subsets are differentially associated with smoking status, COPD severity, and clinical outcomes in NSCLC. Using scRNAseq of lung tissue from non-smokers, smokers, and LUAD, we defined immune signatures linked to smoking exposure and disease state. Subsequent digital cytometry of bulk RNA-seq data from completely resected treatment-naïve tumors revealed that high NK_SR_ cell preservation correlated with better survival following surgery, especially in current smokers. In contrast, in neoadjuvant chemoimmunotherapy-treated NSCLC cohort, tumors enriched with NK_AIR_ cells and depleted of NK_SR_ cells demonstrated superior pathologic responses to chemoimmunotherapy. These findings offer a novel framework for NK cell-based stratification in resectable NSCLC - advancing a biologic clock of immune evasion in which progressive NK_SR_ depletion marks movement toward an immune-suppressed, tumor-permissive state (**Figure 6**).

**Figure 6.**
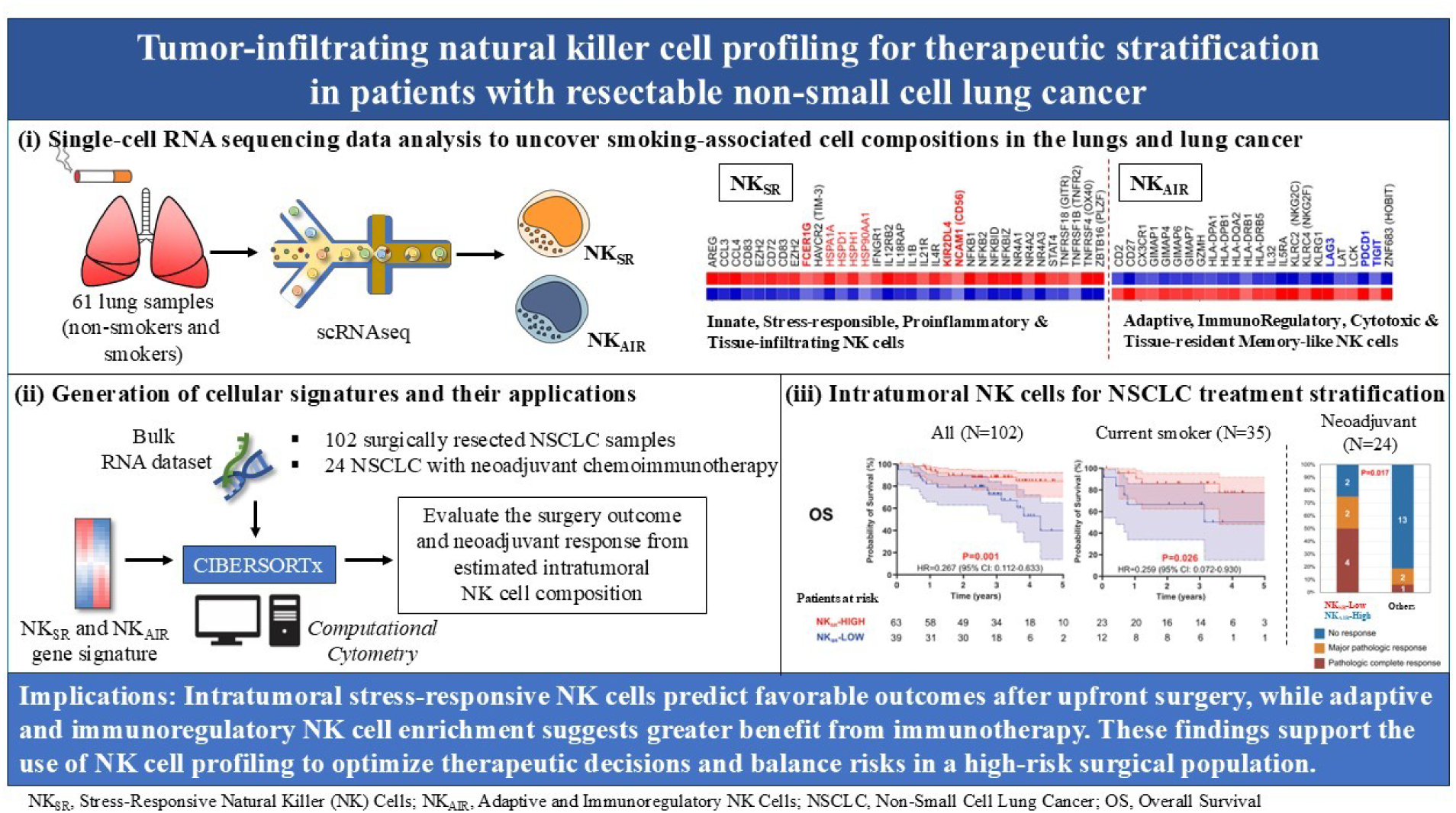
Study design, results, and implications of NK cell profiling for therapeutic guidance in resectable NSCLC.

Neoadjuvant immunotherapy or chemoimmunotherapy offers substantial promise in the treatment of NSCLC by enabling tumor downstaging prior to curative-intent surgery and targeting micro-metastatic disease.^69^ However, successful implementation requires careful multidisciplinary planning to manage treatment-related adverse events^70^ and ensure completion of the planned surgery with minimal postoperative complications.^32, 71^ Current or former smokers often face higher surgical risks due to smoking-related comorbidities and structural lung changes, which are attributable to postoperative complications. Nevertheless, they appear to experience a greater benefit from neoadjuvant chemoimmunotherapy, with approximately 20% achieving a pCR.^25–27^ This observation presents an opportunity for personalized therapeutic strategies, as smokers, who had high surgical risks but a greater likelihood of achieving pCR or MPR, may be prioritized for neoadjuvant chemoimmunotherapy, while patients predicted to derive limited benefit, based on intratumoral NK cell profiling, may proceed directly to upfront surgery (**Figure 7**).

**Figure 7.**
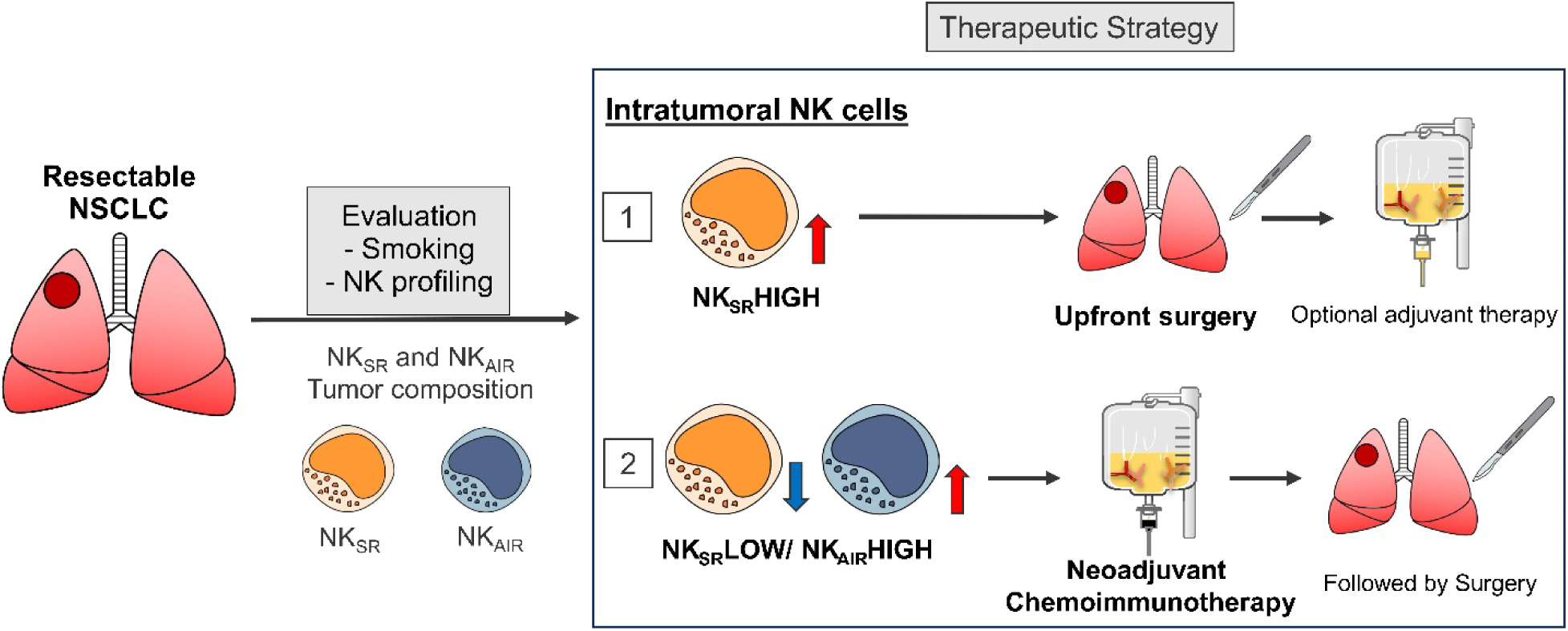
Therapeutic stratification in resectable NSCLC based on intratumoral NK cell composition. A proposed treatment strategy for patients with resectable NSCLC based on intratumoral composition of stress-responsive NK cells (NK_SR_) and adaptive/immunoregulatory NK cells (NK_AIR_). Tumors enriched with NK_SR_ cells (NK_SR_ HIGH) are associated with favorable outcomes and may be effectively treated with upfront surgery (Pathway 1). In contrast, tumors with low NK_SR_ and high NK_AIR_ cell abundance may benefit from neoadjuvant chemoimmunotherapy followed by surgery, resulting in higher rates of complete or major pathologic response (Pathway 2). These findings highlight the potential of NK cell profiling to guide personalized neoadjuvant strategies in NSCLC.

Despite the increasing adoption of neoadjuvant chemoimmunotherapy in smokers with early-stage NSCLC, there remains a critical lack of established biomarkers to guide patient selection. Currently, tumor PD-L1 expression is the only clinically validated and FDA-approved biomarker for predicting immunotherapy response in advanced NSCLC.^72, 73^ However, its clinical utility is hindered by variability in assay platforms, antibodies, and cutoff thresholds.^74^ Another potential biomarker, TMB, faces similar limitations, including challenges in obtaining adequate tissue samples and the limited representation of tumor heterogeneity from a single biopsy.^75, 76^ To address these issues, blood-based TMB is being investigated as a non-invasive alternative.^77–79^ In addition, clearance of circulating tumor DNA (ctDNA) prior to surgery has been associated with improved treatment responses and longer DFS in stage IIIA (N2) NSCLC.^77–79^ While these biomarkers show promise in advanced disease, no validated biomarkers currently exist for predicting response to neoadjuvant chemoimmunotherapy in early-stage NSCLC, highlighting a significant unmet need to characterize the tumor immune microenvironment in this setting. Our study suggests that the intratumoral balance of NK_SR_ and NK_AIR_ cells may serve as a novel predictive immune signature. Tumors with low NK_SR_ and high NK_AIR_ cells were significantly more likely to achieve pCR or MPR following neoadjuvant PD-1 blockade combined with chemotherapy. In contrast, patients with high NK_SR_ infiltration at baseline exhibited favorable outcomes after upfront surgery, suggesting that these tumors may possess sufficient intrinsic immune surveillance, reducing the added benefit of neoadjuvant therapy.

We demonstrated NK cell diversity with NK_SR_ and NK_AIR_ subsets exhibiting opposite correlations with disease severity. NK_SR_ cells are characterized by transcriptional gene profiles with acute inflammation, cytokine responsiveness, stress responses, and rapid activation. NK_SR_ cells are poised for rapid activation under stress and may contribute to the recruitment and maintenance of activated T cells and proinflammatory macrophages in the tumor microenvironment. Conversely, NK_AIR_ cells display a matured, chronically activated, functionally partially exhausted phenotype with tissue-resident memory cell-like features. This subset also exhibited antigen-presenting capabilities with T cell signaling components, representing preserved cytotoxic potential but reduced effector function in chronically inflamed lungs. NK_AIR_ cells expressed multiple immune checkpoint receptors (PDCD1, LAG3, TIGIT), indicating impaired effector function but potential for therapeutic reinvigoration. Notably, NK_AIR_ enrichment in pre-treatment tumors was associated with enhanced response to chemoimmunotherapy, likely due to partial restoration of cytotoxicity following PD-1 blockade and immune remodeling by chemotherapy.^80, 81^

Indeed, neoadjuvant chemoimmunotherapy can substantially remodel the TiME. Recent single-cell studies of stage III NSCLC have shown that neoadjuvant treatment is associated with increased intratumoral NK cell infiltration compared to untreated tumors.^82^ Furthermore, our findings highlight opportunities for therapeutic innovation, such as enhancing NK_SR_ cells via cytokine support (e.g., IL-15 superagonists)^83^, blocking NK-specific checkpoints (e.g., NKG2A)^84^, or adoptive NK cell therapy.^85, 86^ Strategies to prevent stress ligand shedding or recondition the TiME may further potentiate NK cell-mediated antitumor activity.

Several limitations warrant consideration. First, the observational design precludes causal inference. Although we observed strong associations among smoking, NK cell subsets, and clinical outcomes, functional studies are needed to establish mechanistic roles for NK_SR_ and NK_AIR_ cells. Second, NK profiling was performed on resected tumors or bulk RNA-seq, which may not fully reflect the pretreatment immune landscape in small biopsies, although feasibility was demonstrated in a clinical trial.^39^ Future work should develop noninvasive biomarkers, such as circulating or sputum NK phenotypes or soluble NK ligands, to facilitate clinical application. Third, the scRNAseq LUAD cohort and the neoadjuvant-treated cohort were relatively small, underscoring the preliminary nature of these findings. Validation in larger, independent, prospective cohorts will be required. Finally, the influence of co-occurring immune subsets, stromal interactions, and tumor-intrinsic factors was not comprehensively assessed and may affect NK cell function and treatment response.

## CONCLUSIONS

This study explores how smoking may shape the intratumoral immune landscape and examines NK cell heterogeneity in relation to lung immunity and NSCLC outcomes. We describe tumor-infiltrating NK cell profiling as a potential stratification approach for resectable NSCLC. In our data, a higher abundance of stress-responsive NK cells was associated with more favorable surgical outcomes in treatment-naïve NSCLC, whereas tumors enriched for adaptive/immunoregulatory NK cells are more likely to benefit from neoadjuvant chemoimmunotherapy followed by surgery. These observations suggest that intratumoral NK composition could serve as both a prognostic marker and a predictor of response to preoperative therapy. Preoperative tumor NK profiling might help inform individualized decisions to select patients for upfront surgery versus neoadjuvant chemoimmunotherapy. Further studies are needed to validate these NK subsets as clinical biomarkers, develop noninvasive profiling methods, and evaluate strategies aimed at preserving or restoring NK_SR_ or reinvigorating NK_AIR_ function. Together, these efforts have the potential to refine perioperative treatment strategies and improve long-term outcomes for patients with resectable NSCLC by optimizing the balance between therapeutic benefit and risk.

## AUTHOR CONTRIBUTIONS

A.I, H.J.J., F.P., C.I.A, and H.S.L. conceived the study and contributed to the study design, data analysis, interpretation, and writing.

A.I., S.W.K, A.H.S., J.B., and Y.H. contributed to the study design and data analysis. J.S.S., C.L., P.R., G.L., S.M., J.J. contributed to data collection.

F.K., R.T.R., S.S.G., F.P., and B.M.B. provided clinical perspectives and supervisions.

F.P., B.M.B., C.I.A., and H.S.L. contributed to funding acquisition.

## ACKNOWLEDGEMENTS

This work was supported by a Cancer Prevention and Research Institute of Texas grant (Lee: CPRIT RP200443, Amos: RR170048), an NIH R21 (Lee: R21AI159379), a US Department of Defense Impact Award (Lee: CA210552), the Helis Medical Research Foundation (Lee), Samyang Biopharm Research Grant (Lee), NIH R01s (Polverino: R01HL149744 and R01HL171622), an NIH R37 MERIT Award (Burt: R37CA248478), a BCM Department of Surgery Seed Grant (Jang). This project was supported in part by the Genomic and RNA Profiling Core at Baylor College of Medicine with funding from the NIH NCI (P30CA125123) and CPRIT (RP200504) grants.

## Declaration of generative AI and AI-assisted technologies in the writing process

During the preparation of this work, the authors used ChatGPT-5 in order to edit the manuscript. After using this tool/service, the authors reviewed and edited the content as needed and take full responsibility for the content of the publication.

## Conflicts of Interest

- HSL reports research funding from the National Institute of Health, the Department of Defense, the Cancer Prevention Research Institute of Texas, the Helis Medical Research Foundation, and the Dan L. Duncan Comprehensive Cancer Center, as well as investigator-initiated preclinical and clinical research funding from Samyang Biopharm and Momotaro-Gene.
- BMB reports research funding from the National Institute of Health and Cancer Prevention Research Institute of Texas; clinical trial funding from AstraZeneca, Novartis, and Momotaro-Gene Inc.; and has been a consultant in non-small cell lung cancer for AstraZeneca.
- RTR reports research support from the National Institutes of Health, the American Association of Thoracic Surgery, and the DeGregorio Family Foundation; non-remunerated board of director of the Mesothelioma Applied Research Foundation; was retained to provide expert legal opinion; speaker bureau for Merck.
- SSG reports a proctor and speaker honoraria from Intuitive Surgical.
- Other authors have no financial conflict of interest.

## GLOSSARY OF ABBREVIATIONS

COPD: chronic obstructive pulmonary disease
DFS: disease-free survival
GOLD: Global Initiative for Chronic Obstructive Lung Disease
FEV1: forced expiratory volume in 1 second
FVC: forced vital capacity
HLA: human leukocyte antigen
HR: hazard ratio
IFN-γ: interferon-gamma
ILC: innate lymphoid cell
KIR: killer cell immunoglobulin-like receptor
MHC: major histocompatibility complex
MPR: major pathologic response
NF-κB: nuclear factor-kappa B
NK: natural killer
NK_SR_: stress-responsive natural killer
NK_AIR_: adaptive and immunoregulatory natural killer
NSCLC: non-small cell lung cancer
OS: overall survival
PCR: pathological complete response
scRNAseq: single-cell RNA sequencing
TNF-α: tumor necrosis factor-alpha
UMAP: Uniform Manifold Approximation and Projection

