## Supplementary documents for "Tumor-infiltrating natural killer cell profiling for therapeutic stratification in patients with resectable non-small cell lung cancer"

### Table of Contents


| Section | | Page |
|---|---|---|
| **Supplementary Figures** | | |
| **Figure S1** | Single-cell analysis of the lung microenvironment. | 3 |
| **Figure S2** | Integrated single-cell analysis of lung adenocarcinomas. | 4 |
| **Figure S3** | Abundance of NK cell signatures within NK cell populations. | 5 |
| **Figure S4** | Generation of the NK cell abundance score using CIBERSORTx absolute mode adjusted by 22 immune cell phenotypes. | 6 |
| **Supplementary Tables** | | |
| **Table S1** | Patient Characteristics of non-smokers and smokers, relevant to single-cell RNA sequencing data in Figures 1 and 2. | 7 |
| **Table S2** | Patient characteristics of lung adenocarcinoma, relevant to Figure 4. | 8 |
| **Table S3** | ScRNAseq-derived NK cell signatures. | 9 |
| **Table S4** | Univariable regression analyses for overall survival (OS) and disease-free survival (DFS). | 13 |
| **Supplementary references** | | 14 |

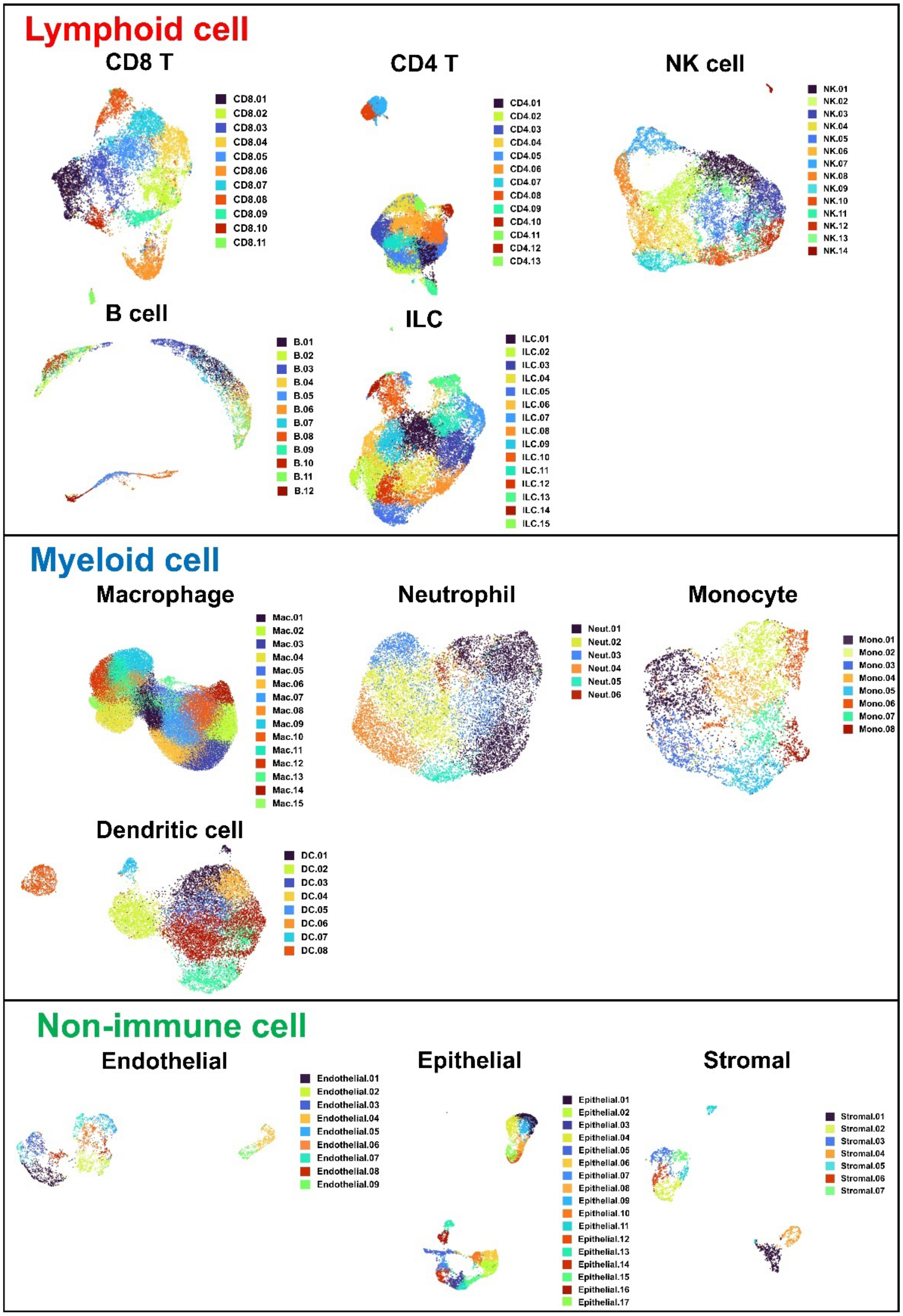


**Supplementary Figure 1**. **Single-cell analysis of the lung microenvironment.**
UMAPs depict unsupervised clustering after lineage stratification into major compartments - lymphoid (CD8 T, CD4 T, NK, B, and innate lymphoid cells (ILC)), myeloid (macrophages, dendritic cells, monocytes, and neutrophils), and non-immune cells (endothelial, epithelial, stromal). Secondary subclustering within each compartment resolved a total of 135 distinct phenotypes. Clusters are color-coded and labeled.

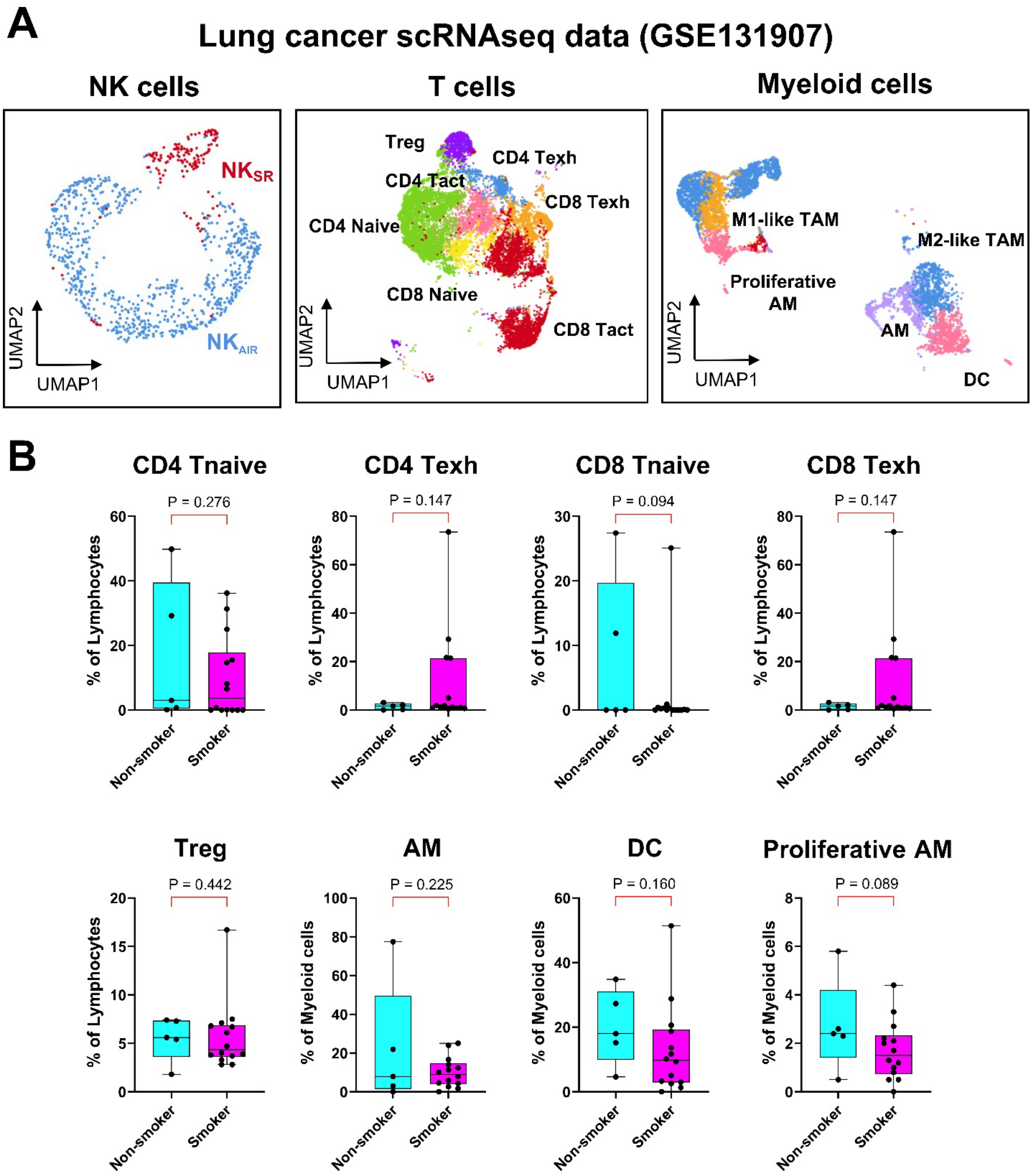


**Supplementary Figure 2. Integrated single-cell analysis of lung adenocarcinomas.**
**A**. Single-cell profiling of 11 LUAD tumors (GSE131907). UMAPs of NK, T, and myeloid compartments are shown. NK cells segregate into $NK_{SR}$ and $NK_{AIR}$ subsets in this independent cohort, with differential gene expression consistent with **Figure 3B**. This dataset was integrated with our BCM scRNA-seq cohort (GSE300685) for the analyses in **Figures 4C, 4E**, and **4G**.
**B**. Integrated single-cell analysis of 19 LUAD tumors (GSE131907+GSE300685). Comparison of immune cell compositions between non-smokers and smokers in LUAD patients.

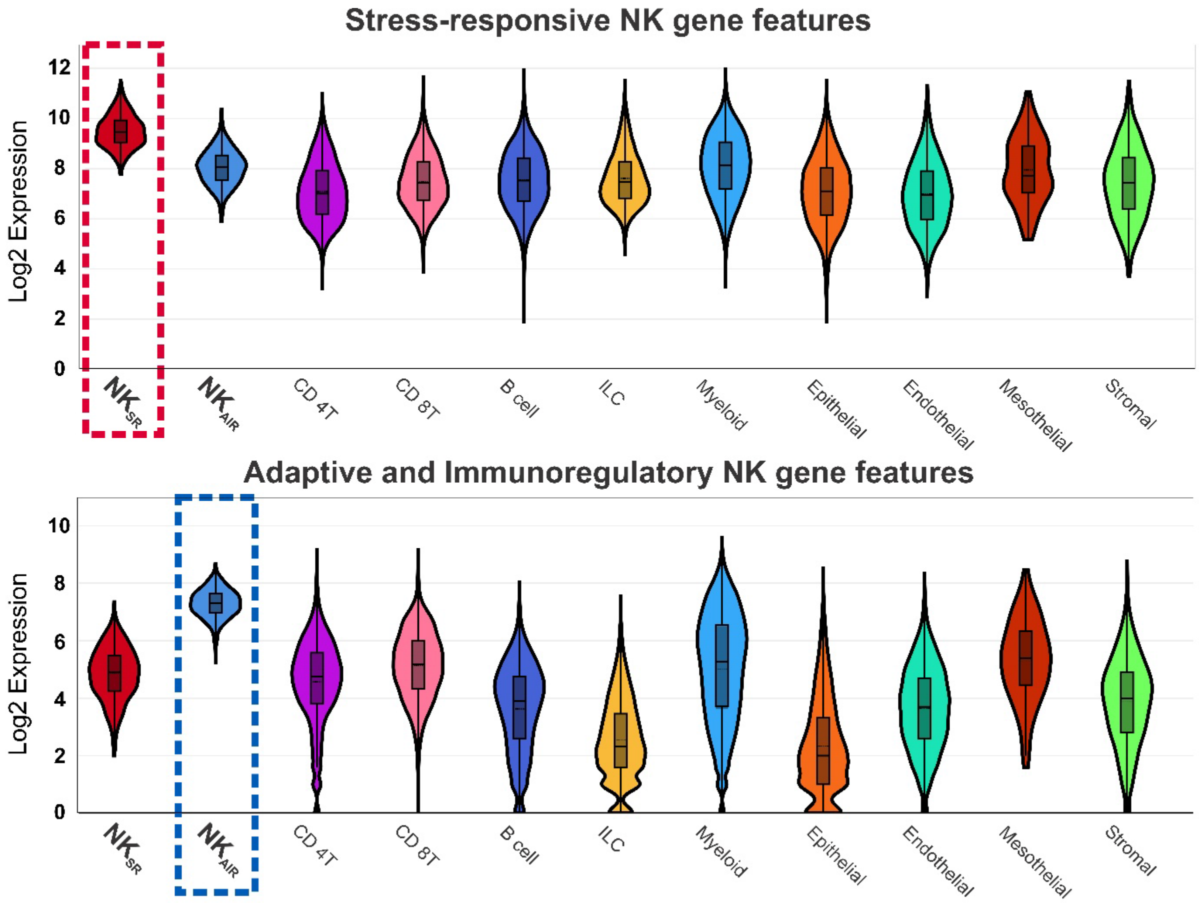


**Supplementary Figure 3. Abundance of NK cell signatures within NK cell populations.**
scRNA-seq-derived NK cell gene signatures were highly enriched in $NK_{SR}$ and $NK_{AIR}$ subsets compared with other cellular populations in the lung microenvironment.

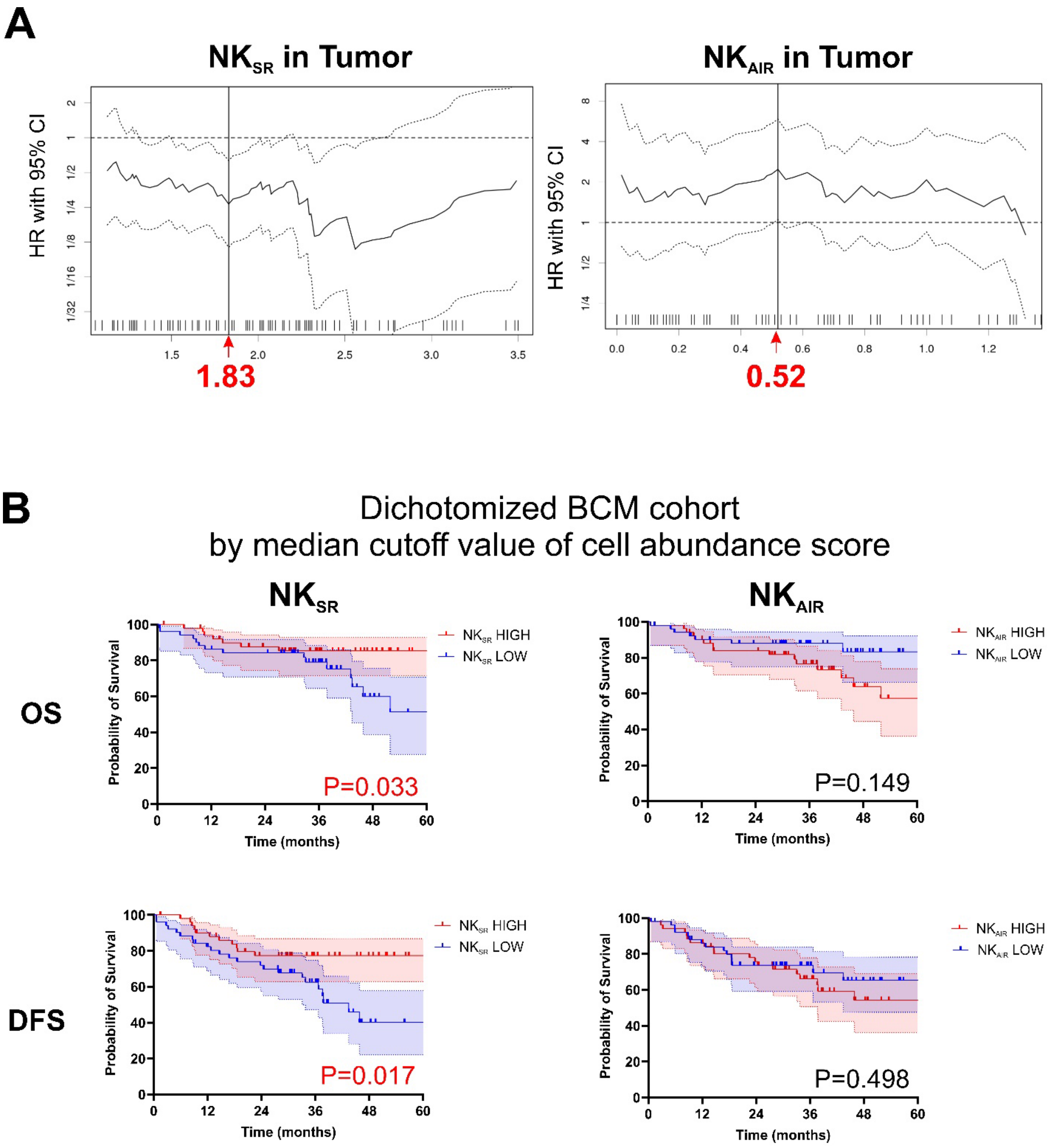


**Supplementary Figure 4. Generation of the NK cell abundance score using CIBERSORTx absolute mode adjusted by 22 immune cell phenotypes.** To quantify NK cell abundance, we utilized the absolute mode of CIBERSORTx (https://cibersortx.stanford.edu/runcibersortx.php), which estimates immune cell fractions adjusted by a reference panel of 22 immune cell phenotypes. Unlike relative cellular proportions, this method provides an absolute abundance score for each immune cell type. **A**. Cutoff values of 1.83 for the NK$_{SR}$ and 0.52 for NK$_{AIR}$ cell abundance score were determined based on the hazard ratio for OS. **B**. Lung cancer patients with high NK$_{SR}$ abundance score demonstrated improved overall survival (OS) and disease-free survival (DFS).

**Supplementary Table 1. Patient Characteristics of non-smokers and smokers, relevant to single-cell RNA sequencing data in Figures 1 and 2.**

| Variables | Non-smoker | Smoker | mild COPD | Severe COPD | *P* value |
|---|---|---|---|---|---|
| Number of patients | 13 | 21 | 15 | 12 | |
| Age, median (range) | 50.2 (23-71) | 71.5 (53-83) | 68.4 (52-78) | 63.3 (35-77) | **<0.001** |
| Sex, n (%) | | | | | 0.469 |
| Male | 8 (61.5) | 10 (47.6) | 8 (53.3) | 9 (75.0) | |
| Female | 5 (38.5) | 11 (52.4) | 7 (46.7) | 3 (25.0) | |
| Smoking Status, n (%) | | | | | **<0.001** |
| Never | 13 (100.0) | 0 (0.0) | 0 (0.0) | 0 (0.0) | |
| Former | 0 (0.0) | 15 (71.4) | 10 (66.7) | 9 (75.0) | |
| Current | 0 (0.0) | 6 (28.6) | 5 (33.3) | 3 (25.0) | |
| PFT, median (range) | | | | | |
| FEV1% predicted | 80 (62-94) | 92 (54-137) | 76 (53-102) | 33 (14-72) | **<0.001** |
| FEV1/FVC | 80 (68-86) | 75 (61-101) | 64 (50-83) | 39 (19-74) | **<0.001** |
| Medications, % | | | | | |
| LABA/LAMA/SABA | 1 (7.7) | 5 (23.8) | 8 (53.3) | 9 (75.0) | |
| Inhaled corticosteroids | 1 (7.7) | 1 (4.8) | 6 (40.0) | 7 (58.3) | |
| Oral corticosteroids | 1 (7.7) | 1 (4.8) | 2 (13.3) | 4 (33.3) | |

*P* values that were statistically significant in the analysis were bolded. *COPD*, chronic obstructive pulmonary disease; *PFT*, pulmonary function test; *FEV1*, forced expiratory volume in 1 second; *FVC*, forced vital capacity; *LABA*, long-acting β adrenoreceptor; *LAMA*, long-acting muscarinic agonist; *SABA*, short-acting β adrenoreceptor.

**Supplementary Table 2. Patient characteristics of lung adenocarcinoma, relevant to Figure 4.**

| Case | GSE No. | Patient ID | Smoking status | Pathologic stage |
|---|---|---|---|---|
| 1 | 131907 | T06 | Former smoker | IA |
| 2 | 131907 | T08 | Non-smoker | IB |
| 3 | 131907 | T09 | Former smoker | IIA |
| 4 | 131907 | T18 | Former smoker | IA |
| 5 | 131907 | T19 | Current smoker | IA |
| 6 | 131907 | T20 | Current smoker | IA |
| 7 | 131907 | T25 | Former smoker | IA |
| 8 | 131907 | T28 | Current smoker | IIIA |
| 9 | 131907 | T30 | Non-smoker | IA |
| 10 | 131907 | T31 | Former smoker | IIIA |
| 11 | 131907 | T34 | Non-smoker | IA |
| 12 | 300685 | LC38 | Former smoker | IIB |
| 13 | 300685 | LC52 | Former smoker | IIIA |
| 14 | 300685 | LC57 | Former smoker | IA |
| 15 | 300685 | LC63 | Former smoker | IB |
| 16 | 300685 | LC71 | Current smoker | IB |
| 17 | 300685 | LC104 | Non-smoker | IB |
| 18 | 300685 | LC115 | Current smoker | IA |
| 19 | 300685 | LC221 | Non-smoker | IB |

**Supplementary Table 3. ScRNAseq-derived NK cell signatures.**

| Gene | $NK_{SR}$ | $NK_{AIR}$ |
|---|---|---|
| AKAP13 | 874.80073 | 382.18026 |
| AKNA | 1136.2998 | 491.81981 |
| ANKRD11 | 401.40609 | 201.1475 |
| AOAH | 1046.5049 | 488.42014 |
| APBA2 | 385.88913 | 126.07132 |
| AREG | 3946.1424 | 215.59613 |
| ARID4B | 538.76939 | 207.09694 |
| ARL6IP5 | 161.27469 | 437.14169 |
| ARPC1B | 216.98314 | 594.66002 |
| ATP1B3 | 906.34341 | 112.75592 |
| AUTS2 | 603.38102 | 129.75431 |
| BHLHE40 | 796.70715 | 171.68364 |
| BTG2 | 1016.7428 | 266.30796 |
| C1orf21 | 382.58223 | 187.83211 |
| CAP1 | 168.90599 | 368.86486 |
| CARD11 | 835.62675 | 186.41557 |
| CBLB | 605.92478 | 299.45481 |
| CCL3 | 1625.4657 | 580.21139 |
| CCL4 | 12362.697 | 4964.6604 |
| CCL4L2 | 1743.4964 | 833.77057 |
| CCL5 | 1507.9438 | 3513.8486 |
| CCNH | 922.36913 | 291.80553 |
| CD247 | 1726.4532 | 771.44317 |
| CD48 | 149.06462 | 375.38091 |
| CD52 | 133.80203 | 973.7239 |
| CD55 | 737.6918 | 201.99742 |
| CD69 | 1752.654 | 759.82761 |
| CD99 | 414.88805 | 951.90931 |
| CDK17 | 685.54462 | 166.01752 |
| CEBPB | 369.86341 | 151.56889 |
| CEBPD | 374.44218 | 83.575371 |
| CEMIP2 | 1534.1446 | 413.91057 |
| CHD1 | 600.83725 | 149.86906 |
| CHST11 | 703.35098 | 222.11217 |
| CLIC1 | 304.74303 | 730.08045 |
| CMIP | 651.2038 | 126.92124 |
| CNOT2 | 463.21958 | 108.78964 |
| CNOT6L | 954.67494 | 203.98057 |
| CORO1A | 388.17851 | 1046.5336 |
| CREM | 899.72962 | 208.23016 |
| CRYBG1 | 474.9209 | 196.6146 |
| DIP2A | 444.90447 | 126.35463 |

| | | |
|---|---|---|
| DNAJA1 | 504.68294 | 187.5488 |
| DNAJB1 | 740.74432 | 301.15464 |
| DNAJB6 | 420.7387 | 153.26873 |
| ELL2 | 915.50097 | 116.72221 |
| EMP3 | 265.0603 | 639.98903 |
| ERN1 | 357.14458 | 146.75269 |
| ESYT2 | 502.90231 | 147.60261 |
| FAM177A1 | 776.61141 | 120.97181 |
| FCER1G | 417.94056 | 134.85382 |
| FNBP1 | 480.51718 | 229.76144 |
| FNDC3B | 391.23103 | 111.33939 |
| FOSL2 | 860.30127 | 143.35301 |
| FTH1 | 3524.6406 | 1711.7369 |
| FYN | 2105.9829 | 792.12454 |
| GIMAP7 | 82.417981 | 395.77896 |
| GMFG | 188.49297 | 383.31348 |
| GNG2 | 1650.9034 | 485.02046 |
| GRASP | 700.55284 | 35.129987 |
| GZMA | 442.36071 | 1004.6043 |
| **GZMH** | 314.15496 | 1292.1602 |
| **HCST** | 529.61184 | 1183.3706 |
| HIPK2 | 434.98379 | 105.10665 |
| HSP90AA1 | 2454.4787 | 917.91255 |
| HSP90AB1 | 1019.5409 | 506.55174 |
| HSPA1A | 787.80397 | 131.17084 |
| HSPD1 | 427.60687 | 127.48785 |
| HSPE1 | 470.5965 | 214.74621 |
| IER2 | 983.92824 | 464.05579 |
| IL2RB | 421.50183 | 194.34815 |
| IL32 | 475.17527 | 2107.7992 |
| INPP5D | 367.57402 | 172.53356 |
| IQGAP2 | 778.39204 | 382.18026 |
| JAK1 | 935.85108 | 452.72354 |
| JARID2 | 374.18781 | 83.008759 |
| JUND | 1764.3553 | 603.44251 |
| KDM6B | 467.79835 | 40.512807 |
| KLF2 | 714.03479 | 1626.745 |
| **KLRC2** | 78.093581 | 602.8759 |
| **KLRK1** | 254.88524 | 526.9498 |
| KMT2E | 605.41603 | 271.12417 |
| LCK | 114.21504 | 403.71154 |
| LYN | 384.10849 | 105.10665 |
| MAP3K8 | 901.51026 | 234.86096 |
| MCTP2 | 633.65182 | 169.98381 |
| METRNL | 1065.0743 | 220.41234 |

| | | |
|---|---|---|
| MT2A | 1158.6849 | 341.10084 |
| MTRNR2L12 | 4342.7153 | 2079.7519 |
| MYL12A | 558.61076 | 1504.3567 |
| NFAT5 | 446.43073 | 105.10665 |
| NFATC2 | 443.63259 | 199.16436 |
| NFE2L2 | 702.0791 | 120.12189 |
| NFKB1 | 1450.4547 | 119.27197 |
| NFKBIA | 2237.7499 | 890.43184 |
| NFKBIZ | 523.25243 | 92.924481 |
| NR4A1 | 378.51221 | 57.227881 |
| NR4A2 | 1325.5559 | 362.06551 |
| NR4A3 | 599.56537 | 61.477477 |
| P2RY8 | 536.22563 | 186.13227 |
| PDE3B | 434.72941 | 114.45576 |
| PDE4D | 698.51782 | 150.71897 |
| PDE7A | 486.36784 | 122.67165 |
| PER1 | 383.09098 | 67.42691 |
| PFN1 | 1760.2853 | 3596.8573 |
| PHF20 | 513.58612 | 187.5488 |
| PIK3R1 | 841.22303 | 349.60003 |
| PITPNC1 | 1018.5234 | 291.80553 |
| PLEK | 185.18608 | 410.7942 |
| PLEKHA2 | 409.80052 | 93.7744 |
| PPP1CA | 143.46834 | 391.24606 |
| PPP1R16B | 537.24313 | 162.90115 |
| PRKCH | 1140.1154 | 540.83181 |
| PRKX | 431.67689 | 109.35625 |
| PSMB9 | 155.16966 | 457.82305 |
| PTGDS | 622.71363 | 181.59937 |
| PTGER4 | 355.61832 | 173.66679 |
| PTP4A2 | 200.70305 | 423.82629 |
| RAB8B | 417.68619 | 161.48462 |
| RAC2 | 441.3432 | 994.40527 |
| RALGAPA1 | 673.84331 | 96.89077 |
| RASA2 | 515.62113 | 220.41234 |
| RASA3 | 664.43138 | 304.27101 |
| RBM38 | 386.1435 | 181.03275 |
| RBM39 | 813.496 | 376.79744 |
| REL | 1056.1712 | 220.41234 |
| RNF125 | 516.89302 | 209.36339 |
| RORA | 599.05662 | 246.75983 |
| RPS4Y1 | 178.06354 | 439.40814 |
| S100A4 | 633.39744 | 1845.4575 |
| S100A6 | 470.34212 | 948.79294 |
| SH3BGRL3 | 801.5403 | 1988.8105 |

| | | |
|---|---|---|
| SIK3 | 364.77588 | 134.28721 |
| SIPA1L1 | 408.52863 | 56.377962 |
| SLA | 750.15625 | 235.71088 |
| SLA2 | 418.44932 | 126.63794 |
| SLC2A3 | 502.90231 | 249.02628 |
| SLC38A1 | 530.62935 | 205.96371 |
| SLC7A5 | 444.14134 | 121.25512 |
| SLC9A3R1 | 176.02853 | 426.65935 |
| SMAP2 | 688.85152 | 308.80391 |
| SMCHD1 | 782.97082 | 336.28463 |
| SORL1 | 388.43289 | 146.46938 |
| SQSTM1 | 691.14091 | 270.84086 |
| SRGN | 2991.4674 | 1432.1136 |
| SSH2 | 667.99265 | 259.50861 |
| STAT4 | 1113.4059 | 276.22369 |
| STK17B | 610.50356 | 302.28787 |
| SYAP1 | 438.79943 | 88.674886 |
| SYTL3 | 2002.1973 | 630.92323 |
| TGFB1 | 1498.7862 | 637.15597 |
| TNFAIP3 | 1800.4768 | 381.89695 |
| TNFRSF1B | 716.57855 | 254.4091 |
| TRAPPC10 | 418.95807 | 185.56566 |
| TSPYL2 | 435.74692 | 137.97019 |
| VAV3 | 717.08731 | 236.8441 |
| VPS37B | 1199.8939 | 173.38348 |
| YES1 | 629.32742 | 147.88591 |
| ZBTB16 | 702.58785 | 49.011997 |
| ZFAND5 | 399.37108 | 103.69012 |
| ZFP36 | 3958.3525 | 1411.7155 |
| ZHX2 | 361.72336 | 56.661269 |
| ZNF331 | 876.58136 | 147.03599 |
| ZSWIM6 | 402.16922 | 61.19417 |

Genes shown in bold indicate NK cell–specific markers.

**Supplementary Table 4. Univariable regression analyses for overall survival (OS) and disease-free survival (DFS).**

| | OS | | | DFS | | |
|---|---|---|---|---|---|---|
| **Variables** | ***P* value** | **HR** | **95% CI** | ***P* value** | **HR** | **95% CI** |
| Sex (female vs. male) | **0.008** | 0.262 | 0.097-0.710 | 0.095 | 0.548 | 0.271-1.109 |
| Age>65yrs | 0.612 | 1.261 | 0.515-3.084 | 0.830 | 0.926 | 0.458-1.872 |
| Race (White) | 0.633 | 0.804 | 0.329-1.968 | 0.804 | 0.908 | 0.424-1.946 |
| Smoking Hx | **0.059** | 1.718 | 0.979-3.016 | 0.312 | 1.256 | 0.808-1.952 |
| Non-smoker | | 1 | | | 1 | |
| Former smoker | 0.067 | 4.103 | 0.906-18.578 | 0.184 | 1.892 | 0.739-4.843 |
| Current smoker | 0.053 | 4.476 | 0.979-20.454 | 0.266 | 1.745 | 0.654-4.652 |
| Smoking amount (Pack-Year) | **0.048** | 1.013 | 1.0-1.025 | 0.124 | 1.009 | 0.998-1.020 |
| FEV1/FVC | 0.250 | 0.191 | 0.011-3.203 | 0.322 | 0.297 | 0.027-3.282 |
| COPD | 0.180 | 1.777 | 0.767-4.116 | 0.456 | 1.315 | 0.641-2.699 |
| Histology (LUSC vs. LUAD) | **0.030** | 2.497 | 1.091-5.712 | 0.137 | 1.725 | 0.840-3.541 |
| PET/CT SUVmax | 0.362 | 1.032 | 0.964-1.106 | 0.955 | 1.002 | 0.943-1.064 |
| Pathological stage | 0.223 | 1.357 | 0.830-2.218 | 0.134 | 1.359 | 0.910-2.031 |
| I | | 1 | | | 1 | |
| II | **0.040** | 2.609 | 1.045-6.518 | **0.012** | 2.656 | 1.240-5.691 |
| III | 0.485 | 1.506 | 0.478-4.747 | 0.390 | 1.506 | 0.592-3.837 |
| High $NK_{SR}$ in the tumor | **0.025** | 0.591 | 0.373-0.937 | **0.034** | 0.671 | 0.463-0.971 |
| High $NK_{AIR}$ in the tumor | 0.309 | 1.429 | 0.718-2.844 | 0.582 | 1.177 | 0.659-2.103 |
| High $NK_{SR}$ in the normal | 0.459 | 0.810 | 0.463-1.416 | 0.967 | 0.991 | 0.628-1.564 |
| High $NK_{AIR}$ in the normal | 0.516 | 1.485 | 0.450-4.899 | 0.403 | 0.647 | 0.233-1.795 |
| Tumor PD-L1 mRNA expression ($log_2$TPM) | 0.401 | 0.882 | 0.657-1.183 | 0.217 | 0.851 | 0.660-1.099 |

*P* values that were statistically significant in the analysis were bolded. *FEV1*, forced expiratory volume in 1 second; *FVC*, forced vital capacity; *LUSC*, lung squamous cell carcinoma; *LUAD*, lung adenocarcinoma; $NK_{SR}$, stress-responsive NK cells; $NK_{AIR}$, adaptive and immunoregulatory NK cells; PD-L1, programmed cell death 1 ligand 1; TPM, transcripts per million.
